# B cells rapidly target antigen and surface-derived MHCII into peripheral degradative compartments

**DOI:** 10.1101/775882

**Authors:** S Hernández-Pérez, M Vainio, E Kuokkanen, V Sustar, P Petrov, S Fórsten, V Paavola, J Rajala, LO Awoniyi, AV Sarapulov, H Vihinen, E Jokitalo, A Bruckbauer, PK Mattila

## Abstract

In order to mount high-affinity antibody responses, B cells internalise specific antigens and process them into peptides loaded onto MHCII for presentation to Th cells. While the biochemical principles of antigen processing and MHCII loading have been well dissected, how the endosomal vesicle system is wired to enable these specific functions remains much less studied. Here, we performed a systematic microscopy-based analysis of antigen trafficking in B cells to reveal its route to the MHCII peptide-loading compartment (MIIC). Surprisingly, we detected fast targeting of internalised antigen into peripheral acidic compartments that possessed the hallmarks of MIIC and also showed degradative capacity. In these vesicles, internalised antigen converged rapidly with membrane-derived MHCII and partially overlapped with Cathepsin-S and H2-M, both required for peptide loading. These early compartments appeared heterogenous and atypical as they contained a mixture of both early and late markers, indicating specialized endosomal route. Together, our data suggests that, in addition to previously-reported perinuclear late endosomal MIICs, antigen processing and peptide loading could start already in these specialized early peripheral acidic vesicles (eMIIC) to support fast peptide-MHCII presentation.

## Introduction

B lymphocytes (B cells) are an essential part of the adaptive immune system, initiating antibody responses against a vast repertoire of different antigens. The presentation of specific antigen-derived peptides loaded onto the major histocompatibility complex (MHC) class II (MHCII) is critical for the ability of B cells to mount a mature antibody response, including class-switch recombination and affinity maturation. In addition, the presentation of peptide-MHCII (pMHCII) complex on the B cell surface enables them to act as antigen-presenting cells (APCs) to CD4+ T lymphocytes (T helper cells, Th cells). T cell receptor (TCR)-pMHCII interaction provides a second activation signal to the B cells and, reciprocally, pMHCII presented on B cells stimulates cognate Th cells to orchestrate other branches of the immune system and to generate CD4+ T cell memory (Whitmire et al., 2009).

Presentation of different antigenic peptides on MHCII is a critical driver of various adaptive immune responses. Other professional APCs, such as dendritic cells (DCs) and macrophages, present peptides from antigens taken up unspecifically by phagocytosis or via receptor-mediated uptake by innate immune receptors, like complement receptors or Fc-receptors. B cells, however, ensure efficient presentation of antigens of given specificity, determined by the B cell antigen receptor (BCR) (Aluvihare et al., 1997; Unanue et al., 2016). Studies on pMHCII loading have largely focused on DCs and macrophages, leaving B cell antigen processing and presentation less understood.

The MHCII peptide-loading compartment (MIIC), where antigen is processed into peptides for loading onto MHCII molecules, is characterized by its main hallmarks, antigen and MHCII. In addition, MIIC contains the key peptide loading chaperone H2-M and the proteolytic enzyme Cathepsin-S (Adler et al., 2017). MIIC has been well characterized by various biochemical fractionation techniques. However, in these assays the information about the heterogeneity, localization and dynamics of the vesicles is typically lost. Therefore, important questions remain about the coordination of antigen processing and MHCII loading and presentation. How the endosomal vesicle machinery of B cells is tuned to enable this highly specific process and how efficient targeting of BCR-bound antigen for processing is coordinated remain unknown. It has been suggested that MIICs are multivesicular and typically contain late endosomal (LE)/lysosomal marker Late Antigen Membrane Protein 1 (LAMP1) (Adler et al., 2017; Lankar et al., 2002; Unanue et al., 2016). Thus, a picture has been outlined where the maturation of MIIC diverts at the stage of multivesicular bodies (MVB) before fusion with end-stage lysosome. However, it is not understood how this process is regulated. To help to decipher the molecular underpinnings of antigen presentation, deeper knowledge on intracellular trafficking of antigen would be required.

In the last 10-15 years, developments in fluorescence microscopy techniques, including improved fluorophores and fluorescent fusion proteins, as well as more sensitive and higher resolution imaging modalities, have significantly increased our general understanding of intracellular vesicle traffic. Microscopy can provide information about the dynamics and heterogeneity of different vesicle carriers that are otherwise challenging to decipher with other techniques. The classical or ubiquitous endolysosomal pathway is delineated as a route from early endosomes (EE) to LE/MVB and, lastly, to lysosomes, with early and late recycling endosomes (RE) sending cargo back to the cell surface. While this general view is relatively well established, new studies continue to reveal dramatic complexity within the endolysosomal system with numerous vesicle sub-populations, transport proteins and vesicle markers as well as vesicle scission and fusion machineries (Chen et al., 2019; Delevoye et al., 2019; Huotari and Helenius, 2011). A group of vital regulators of vesicle traffic are the small GTPases of the Rab protein family that are widely used to define different endolysosomal sub-populations. This family contains more than 60 proteins in humans performing either ubiquitous or specific functions in vesicle traffic (Wandinger-Ness and Zerial, 2014). The appreciation of the role of the Rab proteins has been key in unravelling endosomal network dynamics. However, different carriers vary not only in terms of their Rab identity markers, but also in size, shape, membrane morphology, subcellular localization and acidity. In addition, identification of different cell-type specific variations of vesicular transport systems and diverse specialized endolysosome-related organelles, where MIIC could be included (Delevoye et al., 2019), pose an ongoing challenge for researchers.

In this work, we set up a systematic microscopy approach to follow how antigen, after BCR-mediated internalisation, traffics to MIIC. In accordance with previous studies, we detected and quantified gradual clustering of antigen vesicles towards perinuclear region in 30-60 min. However, already right after internalisation, antigen appeared in heterogenous vesicles that harboured mixed selection of both early and late endosomal markers. Interestingly, these early compartments in the cell periphery possessed hallmarks of MIIC and showed degradative capacity. By specific visualization of membrane-derived MHCII molecules, we found that in these early antigen compartments, MHCII originated largely from the plasma membrane pool, possibly to support fast, first-wave peptide presentation. This study provides the first in-depth imaging of antigen processing pathway in B cells. We found remarkable efficiency in joint targeting of antigen and membrane-derived MHCII into peripheral compartments with hallmarks of MIIC that we name early MIIC (eMIIC). The results increase our understanding of the endolysosomal machinery responsible for MIIC formation and can facilitate future dissections of the regulation of successful antigen presentation.

## Results

### Antigen migrates into the perinuclear area in 30-60 min after activation

To characterize antigen vesicle trafficking in B cells, we first analyse the migration and clustering of antigen in a quantitative manner. We used cultured A20 B cells expressing transgenic D1.3 IgM (A20 D1.3) and activated them with Alexa Fluor-labelled anti-IgM antibodies (AF-αIgM) as surrogate antigen. The localization of the antigen vesicles was imaged in cells fixed at different timepoints and stained for pericentriolar material 1 (PCM1) as a marker for microtubule organizing centre (MTOC) by spinning disc confocal microscopy (SDCM). Well consistent with the literature (Aluvihare et al., 1997; Siemasko et al., 1998; Tsui et al., 2018; Vascotto et al., 2007a) we found that, within 30-60 min, most cells gathered antigen in a cluster that typically localized quite centrally in the cell, in the vicinity of MTOC (Fig. 1A). The same phenomenon was also detected in splenic primary B cells isolated from MD4 mouse strain, selected for their relatively high and homogenous levels of IgM. Primary B cells, however, showed faster kinetics with most cells accumulating antigen in central clusters already in less than 30 min (Fig. 1B). To quantitatively analyse antigen migration, we deconvolved the images to improve the separation of small vesicles and then quantified the total number of vesicles per cell and their mean distance to the MTOC using MATLAB-based 3D analysis (Fig. 1C). By showing a reduction of the vesicle number over time, the analysis clearly demonstrated the fusion, or clustering, of vesicles into bigger entities. At the same time, the average distance to the MTOC decreased, depicting migration of the vesicles closer to the MTOC over time (Fig. 1D, E). Although the vesicle number diminished between 30 and 45 min, the mean distance of the vesicles to the MTOC remained constant. This suggested that the majority of the antigen was trafficked to the perinuclear region already in 30 min, but vesicle fusion events and/or clustering continued at later timepoints (Fig. 1E). The quantification revealed the overall kinetics of the antigen transition from smaller peripheral vesicles into bigger vesicles or vesicle clusters that accumulate close to the MTOC.

**Figure 1.**
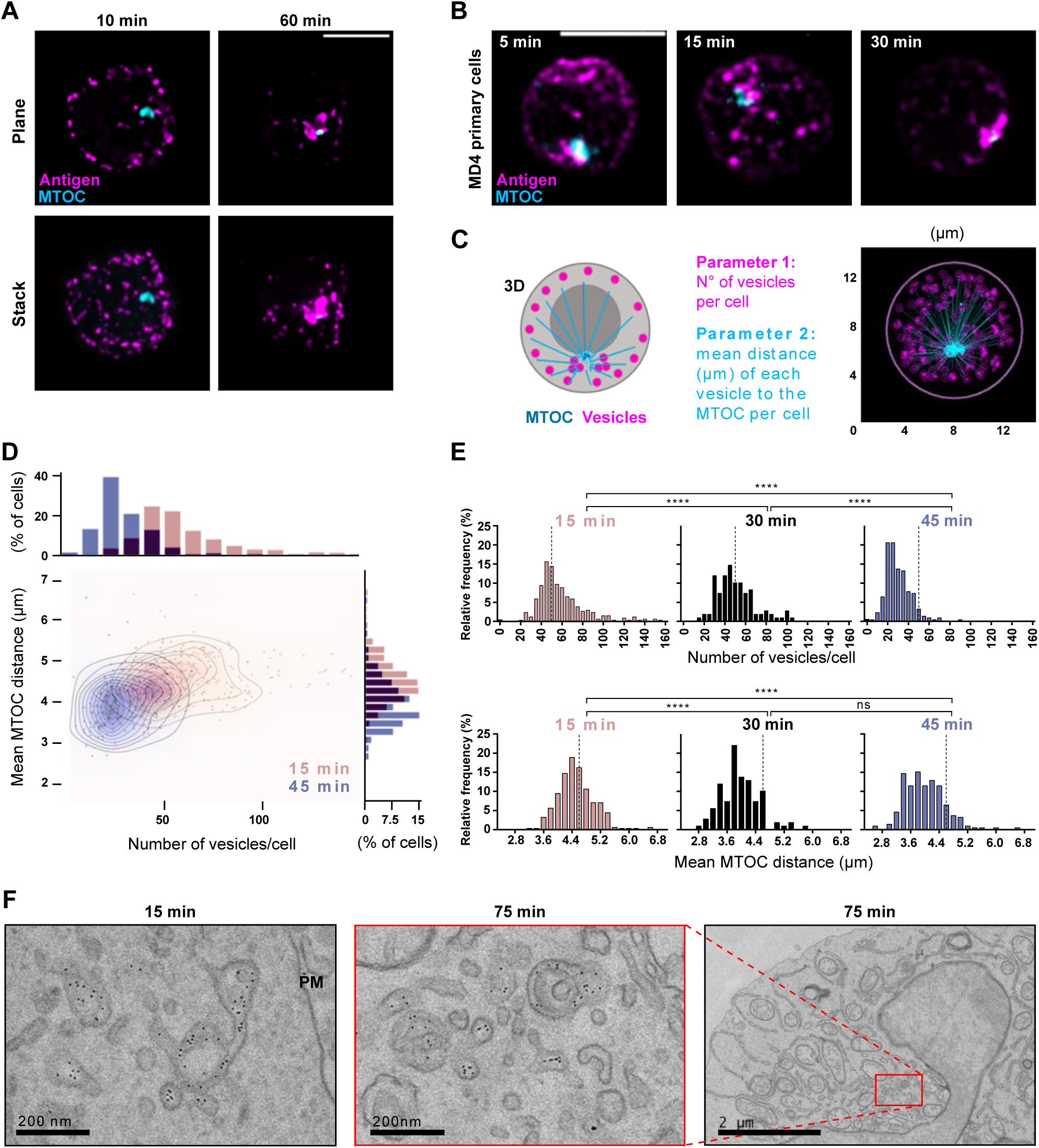
Antigen vesicles traffic to a perinuclear compartment in the vicinity of the MTOC. **A** A20 D1.3 B cells were activated with Alexa Fluor-labelled anti-IgM antibodies (AF-αIgM) (antigen, magenta) for 10/60 min and stained with anti-PCM-1 (MTOC, cyan). Cells were imaged with 3D SDCM and deconvolved. Upper panel, single confocal planes; lower panel, z-projections of 10 µm stacks of representative cells. Scale bar 5 µm. **B** Primary MD4 B cells were activated with AF-αIgM (antigen, magenta) for different timepoints and stained with anti-PCM-1 (MTOC, cyan) and imaged as in A. Z-projections of the whole stacks from representative cells are shown. Scale bar 5 µm. **C** Schematic of the vesicle quantification using a MATLAB-based script. Number of antigen vesicles in one cell (in magenta) and mean distance from all the vesicles to the MTOC (in cyan) is measured in a 3D image. Left, schematic representation; right, example image from the script. **D** Quantification of data in A. 3D images from cells activated for 15 (pink) and 45 (blue) min were analysed as in C. Upper axis, mean number of vesicles per cell; right axis, mean distance of the vesicles to MTOC per cell. The two timepoints were compared using a density plot. **E** Comparison of samples prepared as in A, activated for 15, 30 and 45 min and analysed as in C and D. Dashed line represent the median of the cell population in 15 min. Statistical analysis was done using Student’s t-test. Timepoints 15 and 45 min contain 2 experiments (n>200 cells) and timepoint 30 min one experiment (n>100 cells). **F** A20 D1.3 cells were activated with αIgM conjugated with 6nm colloidal gold particles mixed with AF647-αIgM, for 15 and 75 min and imaged using TEM. PM – Plasma membrane. Scale bars 200 nm and 2 µm.

In order to gain insights into the morphological features of the antigen vesicles, we activated the A20 D1.3 cells with a mixture of AF-αIgM and 6 nm-colloidal gold-αIgM and used transmission electron microscopy (TEM) to visualize antigen-containing membrane structures. We found high heterogeneity in the vesicle morphologies, including multivesicular structures, both after 15 min of activation and after 75 min of activation (Fig. 1F; Fig. S1). As MIICs have earlier been characterized as MVB-like structures (Adler et al., 2017; Lankar et al., 2002; Unanue et al., 2016; Vascotto et al., 2007b), the localization of antigen into multivesicular structures raised a question whether already at 15 min after activation, in the cell periphery, antigen could be processed. At 75 min timepoint, the perinuclear area was, in addition to antigen vesicles, very dense in various other membrane organelles, such as Golgi and mitochondria. Consistent with the literature (Vascotto et al., 2007a), we typically found these vesicle-dense areas at sites of nuclear invaginations (Fig. 1F).

### Antigen colocalisation with early and late endosomal Rab-proteins

Mechanisms of endolysosomal trafficking in various cellular systems are largely governed by Rab-family of small GTPases, which are commonly used to define sub-populations of vesicles with different functions. To reveal the endolysosomal character of the vesicles transporting antigen, we designed a series of colocalisation analyses with the following classical endosomal markers: Rab5 for EEs, Rab7 and Rab9 for LEs and lysosomes, and Rab11 for REs. As antigen-bound BCR is known to form clusters at the cell membrane prior to endocytosis, we first examined the proportion of the dot-like antigen features that was internalised at 10-20 min timepoints, and thus would be expected to colocalise with vesicular markers. At these early timepoints most vesicles still remain in the cell periphery and it is not readily apparent if the antigen is internalised or just clustered at the plasma membrane. To distinguish the internalised antigen from the antigen still on the plasma membrane, we stimulated the cells with biotinylated AF-αIgM and stained with fluorescent streptavidin (Fig. S2A). We detected that in 10-20 min approximately 40-50% of the dotted antigen features in the images represented internalised vesicles, while the rest of the signal originates from antigen that still remains at the cell surface. As expected, at later timepoints (60 min) majority of the antigen pool was detected inside the cells (Fig. S2B, C). This was well consistent with the flow cytometric analysis of antigen internalisation (Fig. S2D). Consistently, we also frequently found non-internalised antigen at the plasma membrane in TEM samples after 15 min of activation (Fig. S1A).

To study the colocalisation of antigen and different vesicle markers, we performed immunofluorescence analysis with SDCM. We expected to see clearly higher colocalisation of antigen with early endosomal Rab5 in the early timepoints, and with LE/MVB markers at later timepoints. To our surprise, we did not detect major differences between the markers. Instead, Rab5, Rab7, Rab9 and Rab11 all showed prominent punctate pattern of vesicles very close to the plasma membrane that partially overlapped with antigen and partially located just underneath the antigen signal (Fig. 2A, C; Fig. S3A). As a negative control, we used Golgi-specific transport protein Rab6 and, as expected, it showed no notable colocalisation with antigen. We quantified the colocalisation using Manderś overlap coefficient, split for antigen channel (M2) (Manders, E. M.M. Verbeek, F. J. Aten, 1993), using Huygens software with automated thresholding. M2 measures the portion of antigen that overlaps with the signal from different Rab-proteins. The analysis supported partial colocalisation of antigen with Rab5, Rab7, Rab9 and Rab11, already at early timepoints after activation (Fig. 2C).

**Figure 2.**
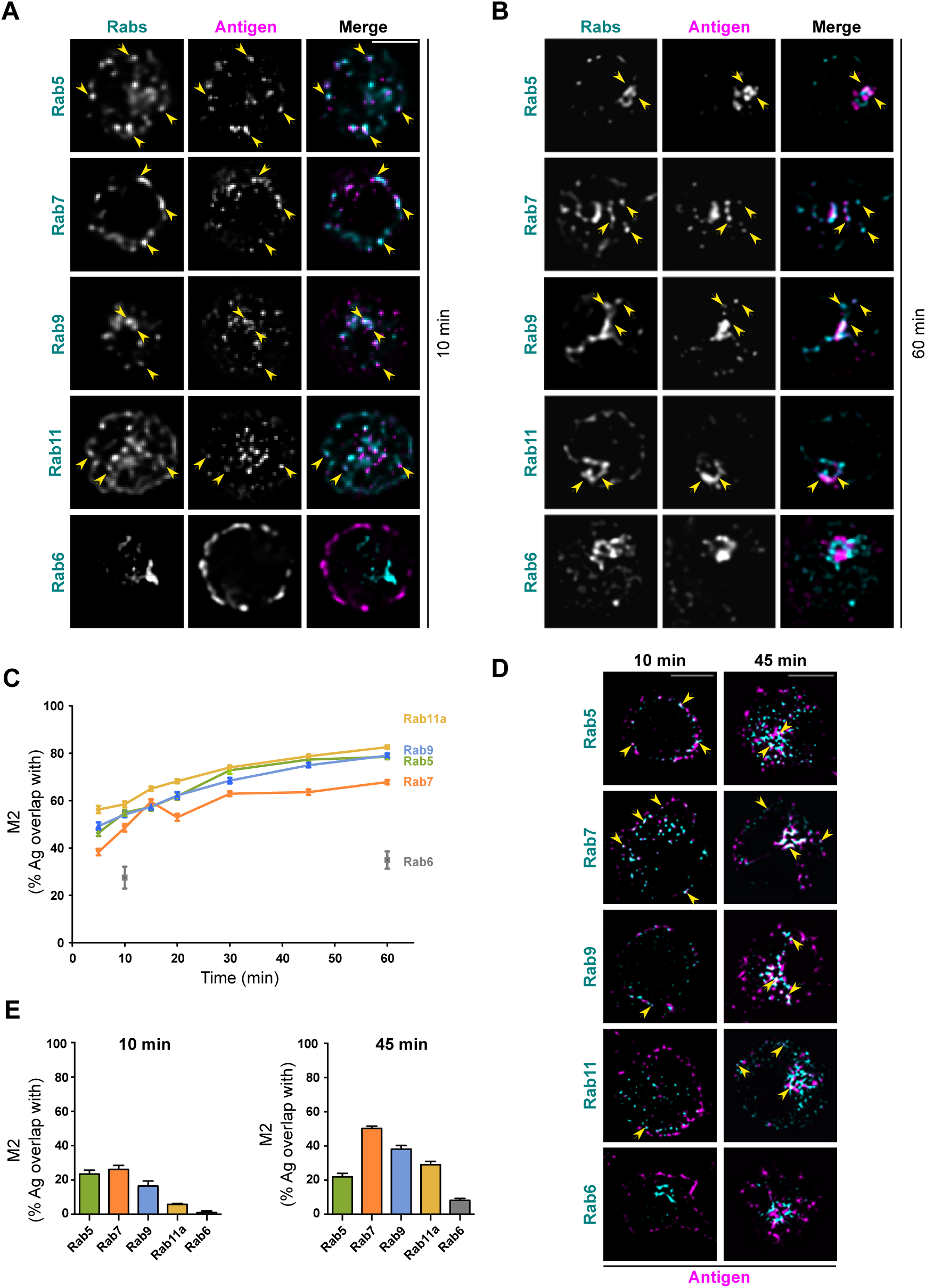
Colocalisation analysis of antigen with different Rab-proteins. **A-B** SDCM imaging of A20 D1.3 cells activated with AF647-αIgM (antigen, magenta) for 10 min **(A)** or 60 min **(B)** and immunostained for different Rab-proteins: Rab5, Rab7, Rab9, Rab11 and Rab6 (cyan). Single confocal planes from deconvolved representative cells are shown and examples of colocalising vesicles are pointed with yellow arrow-heads. For clear representation, single confocal planes close to the bottom of the cell are shown for Rab5, Rab7, Rab9 and Rab11. For Rab6, a confocal plane from the middle of the cell, were Golgi is typically located, was selected. See Figure S3A-B for Z-projections. Scale bar 5 µm. **C** Quantification of the data in A and B with additional timepoints. Antigen colocalisation with different Rab-proteins was measured from deconvolved images analysing Manderś overlap coefficients using Huygens. Data from three independent experiments (>80 cells/timepoint) as mean ±SEM. **D** Samples were prepared with cells activated for 10 or 45 min as in A-B and imaged with iterative imaging of a single plane with SDCM (20-25 frames/plane) and post-processed to obtain SRRF super-resolution image (antigen, magenta; Rabs, cyan). Examples of colocalising vesicles are pointed with yellow arrowheads. Scale bar 5 µm. **E** Quantification of the SRRF data in D analysing Manderś overlap coefficients with ImageJ. Data shown as mean ±SEM. 45 min timepoint, two independent experiments; 10 min, one experiment (>25 cells/timepoint).

At 60 min timepoint, when most of the antigen was clustered in the perinuclear region, we found enhanced colocalisation with LE/MVB markers Rab7 and Rab9, as expected (Fig. 2B, Fig. S3B). Rab11, involved in slow recycling, also localized to the antigen cluster as well as the EE marker Rab5. The negative control, Rab6, was found in the perinuclear region close to antigen but with very limited overlap in signals. This suggested translocation of the antigen close to the Golgi apparatus, supporting the above observed localization close to the MTOC (Fig. 1). The quantification suggested significant overlap of antigen with all the studied Rab-proteins except Rab6, with an increasing trend over time (Fig. 2C). We also analysed the possible effect of antigen uptake in the distribution of Rab proteins by comparing the intensity of different Rab+ compartments in the vicinity of MTOC to the intensity throughout the cell before or 10 and 45 min after activation (Fig. S4A, B). The change in the distribution was most clear in the case of Rab7, which concentrated significantly closer to the MTOC at late time points after activation, accompanied by a decrease in the number of Rab7 vesicles (Fig. S4C, D). However, interestingly Rab9 distribution did not show any alteration, indicating functional divergence between these two LE/MVB markers. On the other hand, Rab5 and Rab11 showed rather increased dispersion away from MTOC after 10 minutes, which could be explained by the activation of the endocytic and exocytic machineries in the case of Rab5 and Rab11, respectively.

Majority of the vesicles were found very close to each other, both at the early and late time points, at the vicinity of the plasma membrane or in the perinuclear region, respectively, leading to overestimation of the signal overlap. The antigen vesicles detected by spinning disk confocal microscope, after deconvolution and the analysis by MATLAB script (as in Fig 1), range between 200nm and 1μm in diameter, with a large majority of vesicles falling in between 400-500 nm (data not shown). Based on EM micrographs, the actual size of the antigen-containing unilamellar vesicles was, however, found to be ≈120 nm and multilamellar vesicles ≈290 nm (Fig 1F, Fig S1A, C), suggesting that the apparent vesicle sizes in fluorescence microscopy are affected by the limited resolution in optical microscopy. Small vesicles located close together are not resolved individually, but appear as one or several larger vesicles. In order to improve the resolution of our data and to better separate different vesicles, with a method still suitable for relatively large sample numbers and quantification, we employed super-resolution radial fluctuations (SRRF), an imaging method based on post-processing analysis of signal fluctuations (Gustafsson et al., 2016). Here, we analysed samples activated for 10 or 45 min in order to resolve the nature of the antigen vesicles in the perinuclear region. Super-resolution SRRF images were obtained by taking 20-50 repetitive images of the same field of view with SDCM and post-processing the data using SRRF plugin in ImageJ. In this way, we could improve the separation of the vesicles significantly and now detected more distinct differences in the localization of Rab-proteins with respect to antigen, especially in later timepoints. In 45 min, Rab7 and Rab9 showed clear colocalisation with antigen, as expected for their late endosomal nature, while Rab5 and Rab11 appeared more scattered and only partially colocalised with antigen, often marking vesicles or membrane domains adjacent to it (Fig. 2D and Fig. S5). We analysed SRRF images for Manderś overlap coefficient (M2) and detected a marked colocalisation of antigen signal with LE markers Rab7 and Rab9 in 45 min. We also detected overlap with Rab11, and, to some extent, with Rab5. However, in 10 min the analysis showed close to equal colocalisation of Rab5, Rab7 and Rab9, and a modest level of colocalisation with Rab11. Colocalisation of antigen with Rab6 remained low, confirming the specificity of the analysis (Fig. 2E). The presence of Rab11 in the antigen vesicles could suggest fission and recycling to some extent already at early timepoints, but with increasing efficiency towards later timepoints.

All together, these results point towards a previously unnoticed heterogeneity in the antigen-containing endosomes. Early association of antigen with classical LE/MVB markers Rab7 and Rab9 raises the possibility that antigen vesicles deviate from classical steps of EE to LE conversion during their maturation into MIIC.

### Antigen trafficking involves atypical vesicles that share both early and late endosomal character

To better define antigen transport vesicles, we asked how other typical EE and LE markers, Early Endosome Antigen 1 (EEA1) and LAMP1, respectively, correlated with antigen at different timepoints. Consistent with the data on different Rab-proteins, we detected partial colocalisation of antigen with both EEA1 and LAMP1 already at early timepoints (Fig. 3A, Fig. S3C). The colocalisation became more prominent as antigen trafficked to the perinuclear region (Fig. 3B, Fig. S3D). Manderś overlap coefficient also showed continuous, or perhaps even increasing, overlap with antigen for both markers (Fig. 3C). However, Pearsońs correlation coefficients for EEA1 and LAMP1 crossed over time indicating that significantly higher proportion of LAMP1 compared to EEA1 colocalised with antigen at later timepoints. This is consistent with a high proportion of EEA1 endosomes remaining in the cell periphery, while some coalesce in the central cluster together with antigen, as shown by the M2. On the other hand, increasing proportion of LAMP1-positive vesicles accumulated in the perinuclear region with antigen over time.

**Figure 3.**
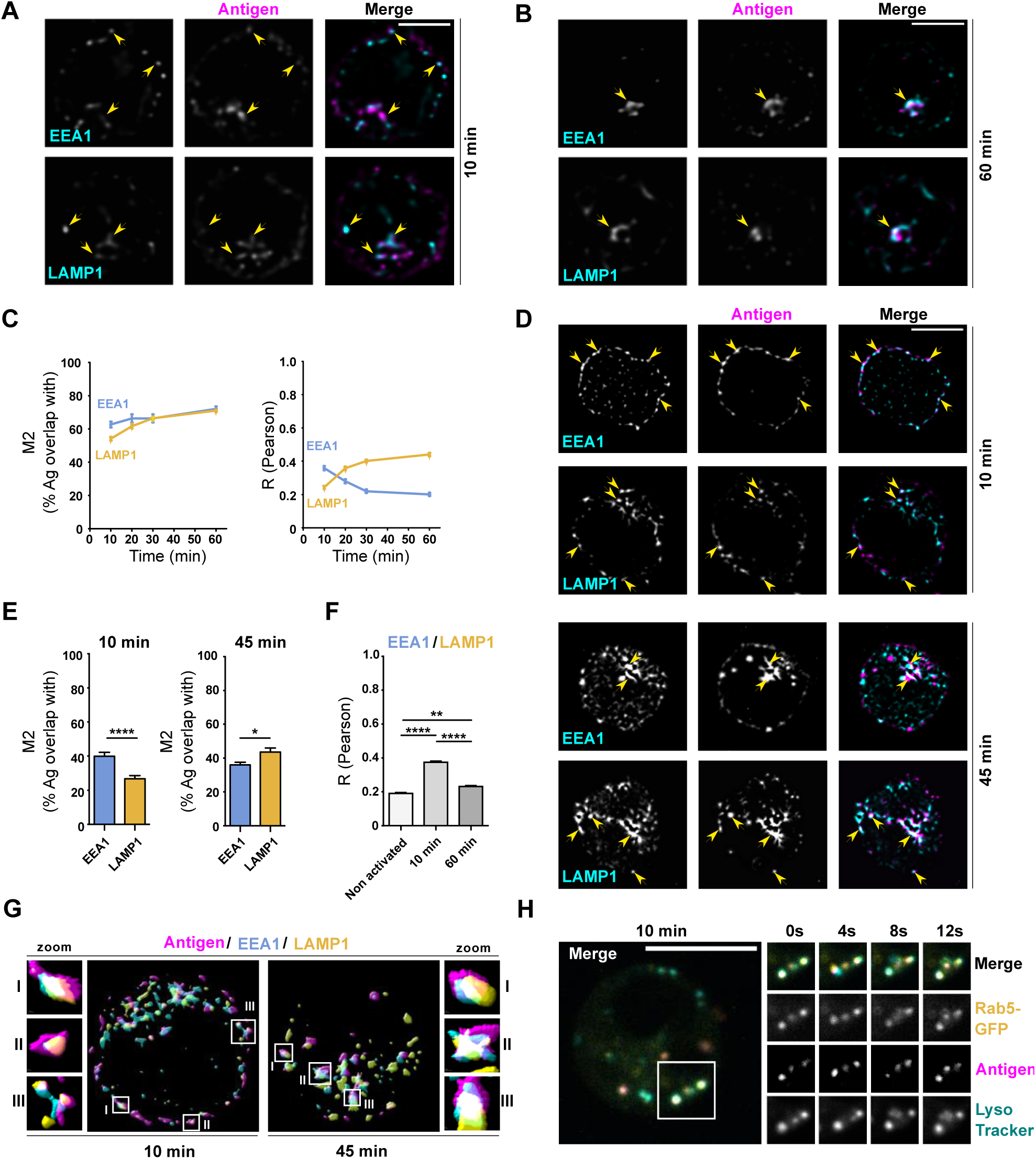
Antigen colocalises with both EEA1 and LAMP1 while trafficking to the perinuclear region. **A-B** SDCM imaging of A20 D1.3 cells activated with AF647-αIgM (antigen, magenta) for 10 min **(A)** or 60 min **(B)** and immunostained for EEA1 or LAMP1 (cyan). Single confocal planes from deconvolved representative cells are shown and examples of colocalising vesicles are pointed with yellow arrow-heads. See Figure S3C-D for Z-projections. Scale bar 5 µm. **C** Quantification of the data in A and B with additional timepoints. Antigen colocalisation with EEA1 and LAMP1 was measured from deconvolved images analysing Manderś overlap coefficients and Pearsońs correlation coefficients using Huygens. Data from two independent experiments (>40 cells/timepoint) shown as mean ±SEM. **D** Samples were prepared with cells activated for 10 or 45 min as in A-B and imaged with iterative imaging of a single plane with SDCM (20-25 frames/plane) and post-processed to obtain SRRF super-resolution image (antigen, magenta; EEA1/LAMP1, cyan). Examples of colocalising vesicles are pointed with yellow arrowheads. Scale bar 5 µm. **E** Quantification of the SRRF data in D analysing Manderś overlap coefficients with ImageJ. Data from three independent experiments (>30 cells/timepoint) shown as mean ±SEM. **F** Quantification of EEA1/LAMP1 colocalisation by analysing Pearsońs correlation coefficient with Huygens. Data from three independent experiments (>30 cells/timepoint) shown as mean ±SEM. **G** Surface reconstruction using Huygens rendering tool of SRRF images from samples prepared as in D (antigen, magenta) and immunostained for EEA1 (cyan) and LAMP1 (yellow). Three selected example vesicles are highlighted by zoom-in. **H** A20 D1.3 cells were transfected with GFP-Rab5 (yellow), loaded with LysoTracker (LT; cyan) and activated with RRx-αIgM (antigen, magenta). Live-imaging was performed with SDCM (ORCA camera) on a single plane. On the left, a merge image of a representative cell after 10 min of activation is shown. On the right, the region in the white square is followed in split channels as a timelapse for 12 s, starting 10 min after activation. See Movie S1.

As a complementary approach, we again turned to SRRF super-resolution analysis in order to achieve higher accuracy. We examined the colocalisation of antigen with EEA1 and LAMP1 at 10 and 45 min after activation. SRRF analysis confirmed higher colocalisation of antigen with EEA1 compared to LAMP1 in the early timepoints (10 min), and vice versa after 45 min (Fig. 3D, E). Nevertheless, EEA1 colocalisation with antigen was also detected both in some remaining peripheral vesicles and in the perinuclear antigen vesicle cluster, raising a possibility that also EEA1 could indeed localize to the MIIC. Together, this data revealed surprising localization of antigen with not only early, but also late endosomal carriers shortly after internalisation. At later timepoints, close to the MTOC, preference for LE/MVB markers was notable, yet also EE markers were found to overlap with antigen.

As previous shown (Fig. S2B-C), in 15 min approximately half of the dots with antigen signal should represent vesicles inside the cell, and the rest should originate from antigen-BCR clusters still at the plasma membrane. Therefore, M2 values for one type of vesicle marker should not be considerably higher than 50% in the early timepoints. Our observation that antigen showed an overlap of 40-60% with both early and late endosomal markers can simply reflect technical challenges to resolve small vesicles close to each other, causing adjacent vesicles to appear as colocalised. Additionally, it could point towards mixed vesicle identities that would simultaneously possess both types of markers. To test for these two non-exclusive scenarios, we next performed SRRF super-resolution analysis on cells activated either for 10 or 45 min and stained for LAMP1 and EEA1, and asked if they colocalised in the same antigen vesicles. We found vesicles, where antigen only colocalised with either EEA1 or LAMP1, but we also found several prominent vesicles that clearly contained both markers simultaneously (Fig. 3G). To investigate if this atypical colocalisation was triggered by antigen uptake, we next analysed non-activated cells. Interestingly, we found vesicles with clear EEA1/LAMP1 colocalisation already in resting cells. The quantification by Pearson’s coefficient showed lower colocalisation in resting cells as compared to cells activated for 10 min, but comparable colocalisation as in the cells activated for 45 min (Fig 3F; Fig. S6A). In order to seek for further confirmation to these findings, we also analysed the colocalisation between other pairs of early and late endosomal markers, namely Rab5/LAMP1, Rab7/EEA1 and Rab9/EEA1, before and after activation. In all cases, already at the resting state, we detected some vesicles with colocalisation, but very low overall level of correlation, which again, however, increased in 10 min after antigen stimulation (Fig. S6B).

Next, we asked if the vesicles that share both early and late endosomal markers were in the transition state of their maturation or if they represented a special compartment. To investigate this, we performed live imaging of A20 D1.3 B cells transfected with green fluorescent protein (GFP)-fused Rab5 and loaded with LysoTracker, a fluorescent tracer that labels low pH compartments, such as LE/MVBs and lysosomes. We followed antigen vesicles at early timepoints after internalisation by SDCM. We detected several antigen vesicles that contained both Rab5 and LysoTracker (Fig. 3H; Movie S1). Joint movement of the markers implied physical colocalisation, and indicated that antigen, indeed, traffics in atypical vesicles that share both early and late endosomal features. Interestingly, we detected double positive vesicles also before cell activation (Fig. S6C).

### Antigen enters degradative compartments shortly after internalisation

As the primary purpose of antigen uptake by B cells is to degrade it for loading the resulting peptides onto MHCII complexes, we next asked the question where and when does the antigen degradation start. We linked a fluorescent probe for proteolysis, DQ-OVA, to αIgM or specific HEL antigen recognized by the D1.3 BCR (Fig. 4A). Fluorescent DQ moieties quench each other when the probe remains intact. However, upon proteolysis the quenching ceases and the fluorescence can be detected. We first analysed the increase in DQ fluorescence by flow cytometry. Already at 15-20 min we detected a clear signal that constantly increased through the analysis period, 45 min, suggesting that the proteolysis starts relatively fast after antigen uptake (Fig. 4B). We detected brighter DQ-Ova signal at later timepoints when linked to HEL, despite having similar internalisation rate to anti-IgM (see Fig. S2D). Due to the brighter signal of DQ-Ova conjugated to HEL, we performed the microscopy analysis using this probe. Fluorescence signal from antigen-linked DQ moieties was also visible by microscopy at 20 min after activation. The DQ-signal overlapped well with EEA1 and was also found to colocalise with CatS, an enzyme essential for preparing the MHCII for peptide loading (Fig. 4C).

**Figure 4.**
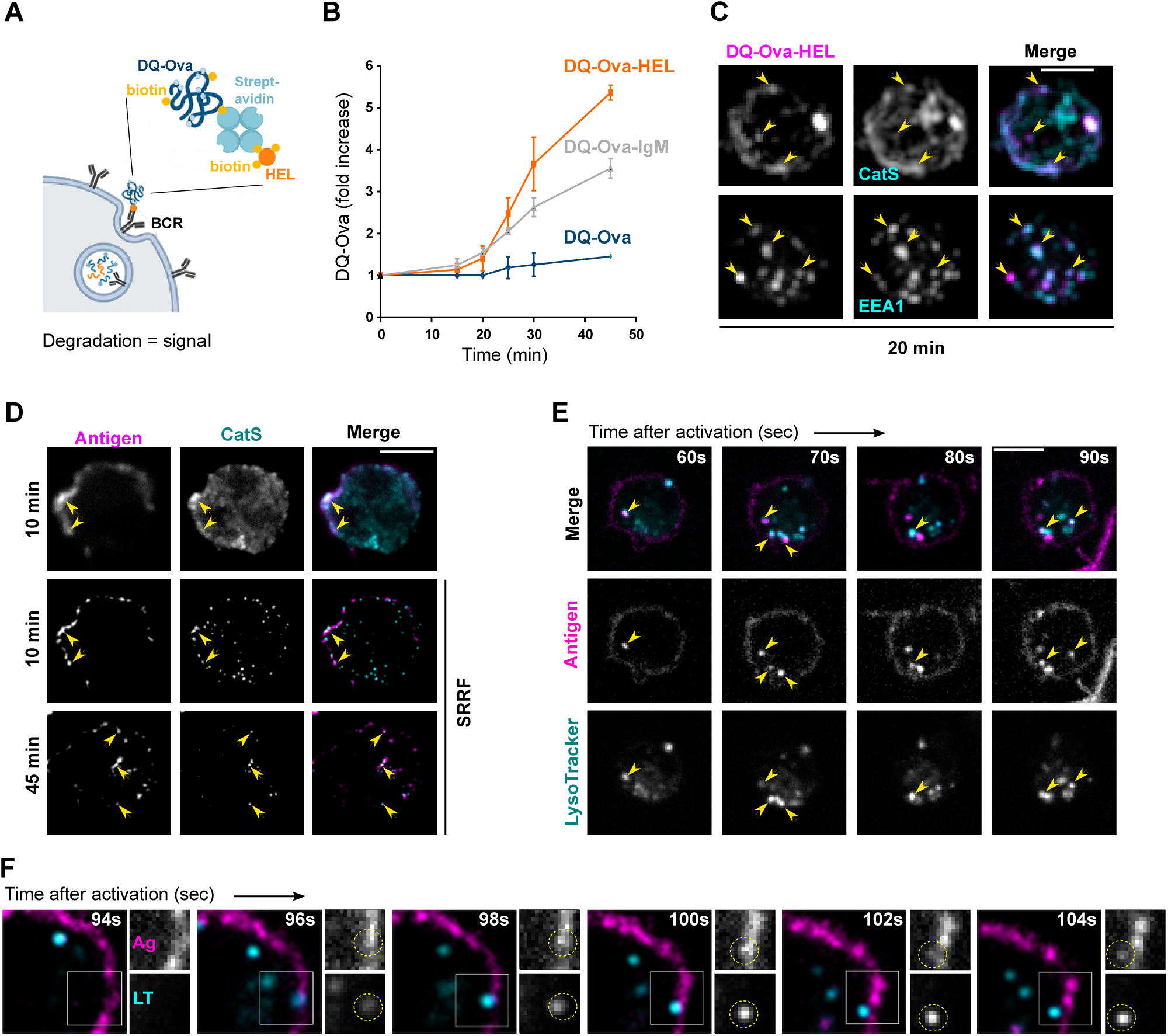
Internalised antigen incorporates into vesicles with low pH and capability to degrade cargo. **A** Schematic view of DQ-Ova-antigen (HEL) sandwich to probe proteolysis of antigen internalised by the BCR. **B** DQ-Ova and DQ-Ova-antigen (αIgM or HEL) degradation assessed by flow cytometry. Cells were labelled as in A, washed, and incubated for different timepoints at 37°C and fluorescence of DQ-OVA was acquired immediately. Results are shown as fold increase (mean ±SD of the DQ-OVA intensity, normalized to the intensity at time zero). N>2 independent experiments. **C** A20 D1.3 cells activated with DQ-Ova-HEL (magenta) as in B, for 20 min, were immunostained for EEA1 or CatS (cyan). Images were acquired using SDCM with EVOLVE (EMCCD) camera. Z-projections of representative cells (n = 3 independent experiments) are shown with examples of colocalising vesicles pointed with yellow arrow-heads. Scale bar 5 µm. **D** A20 D1.3 cells activated with AF647-αIgM (antigen, magenta) for 10 or 45 min and immunostained for CatS (cyan) were imaged with conventional SDCM (upper panel, single plane) or with iterative imaging to obtain SRRF super-resolution image (20-25 frames/plane) (middle and bottom panels). Examples of colocalising vesicles are pointed with yellow arrowheads. Scale bar 5 µm. **E-F** A20 D1.3 were loaded with LysoTracker (cyan) and activated with AF488 F(ab’)2-αIgM (antigen, magenta). Live-imaging was performed with SDCM with EVOLVE (EMCCD) camera every 2s (E) or 500 ms (F), starting as soon as possible after transition of the cells to 37°C under the microscope. **(E)** A timelapse from a representative cell is shown and examples of colocalising vesicles are pointed with yellow arrowheads. Scale bar 5 µm. See Movie S2. **(F)** A timelapse of an example movie highlighting a probable fusion event between an internalising antigen vesicle and a LT vesicle (dashed yellow circle). A white square in the merge image (left) depicts the region of the split channel insets. See Movie S3.

To investigate the level of antigen colocalisation with CatS in a more comprehensive way, we performed immunofluorescence analysis in cells activated for 10 or 45 min. Conventional SDCM imaging suggested partial colocalisation of CatS with antigen both at 10 and 45 min timepoints (Fig. 4D, upper panel). In order to resolve the vesicles better, we performed SRRF analysis, and could more unambiguously detect antigen vesicles that clearly contained CatS already 10 min after activation (Fig. 4D, middle panel). Interestingly, the colocalisation level remained roughly similar, although low, in the later timepoints in the perinuclear region (Fig. 4D, bottom panel; Fig. S6D). Proteolytic activity typically requires acidic pH of the vesicles. To examine the pH of the antigen vesicles, we used live imaging with LysoTracker, as its accumulation is based on acidic pH. In line with our data above (Fig. 3H), we found strong colocalisation of antigen with LysoTracker already in the very early timepoints (1-5 min after activation) (Fig. 4E; Movie S2). Notably, we also detected antigen fusing with LysoTracker positive vesicles immediately after internalisation, indicating very fast and efficient targeting of antigen to acidic vesicles (Fig. 4F; Movie S3). Curiously, LysoTracker positive vesicles appeared to hover beneath the plasma membrane ready to catch the internalised antigen.

In addition to LysoTracker, we indirectly studied vesicle pH by analysing the fluorescent decay of FITC coupled to α-IgM. While AlexaFluor fluorophores are highly stable at acidic pH, FITC fluorescence is pH-sensitive. A20 D1.3 cells were activated using both AF647-conjugated α-IgM, as a control, and FITC-conjugated α-IgM as pH probe, and fluorescence intensities were followed over time by flow cytometry. In agreement with our results using DQ-Ova and fast colocalisation with lysotracker+ compartments, we observed a decay in FITC signal already at 5 minutes after internalization that continued to further decrease through the experiment (Fig S6E). In contrast, AF647 signal remained constant, indicating high stability of the fluorophore.

### Antigen colocalises with plasma membrane derived MHCII rapidly after internalisation

The data above suggests that antigen processing could be initiated already in the peripheral antigen vesicles shortly after internalisation. To ask if these early vesicles might represent MIIC, we asked whether they also contain MHCII. We activated the cells for 10 or 60 min with fluorescent antigen and performed immunofluorescence staining of total MHCII. As expected, we found MHCII to strongly colocalise with antigen in the perinuclear antigen cluster at 30 min timepoint. Interestingly, after 10 min of activation, we also detected MHCII in various intracellular vesicles including those containing antigen (Fig. S6F). Due to the high signal originating from the plasma membrane-resident MHCII and several internal MHCII-positive structures, we decided to analyse the samples with a super-resolution technique structured illumination microscopy (SIM). SIM significantly improved the resolution and clarity of the imaging (x-y-z), and we could detect strong colocalisation of antigen and MHCII in clearly defined vesicles already at 15 min after cell activation. We detected extremely high M1 Manderś overlap coefficients in the areas with peripheral antigen vesicles indicating that almost all internalized antigen overlapped with MHCII. A significant proportion of MHCII was also found with antigen in these regions, further supported by Pearson’s correlation coefficients (Fig. 5A). To investigate if the MHCII in the early antigen vesicles was newly synthesized from the trans-Golgi network, or originated from the plasma membrane pool, we prelabelled the surface MHCII prior to cell activation (Fig. 5B). Interestingly, we saw a strong localization of surface-derived MHCII (sMHCII) to early antigen vesicles (Fig. 5C). We then proceeded to verify the colocalisation by performing live imaging of cells labelled with fluorescent anti-MHCII prior to activation with fluorescent antigen. The movies revealed very high level of sMHCII in the antigen vesicles (Fig. 5D, Movie S4). Finally, to further prove that the early antigen vesicles could function as MIIC, we stained the cells for H2-M, a molecule of MHC family that functions as a key chaperone in peptide loading to MHCII (Mellins and Stern, 2014). Notably, SRRF super-resolution imaging of immunofluorescence samples showed clear colocalisation of antigen vesicles and H2-M already at 15 min after activation further supporting classification of these vesicles as early MIICs (eMIICs) (Fig. 5E; Fig. S6D).

**Figure 5.**
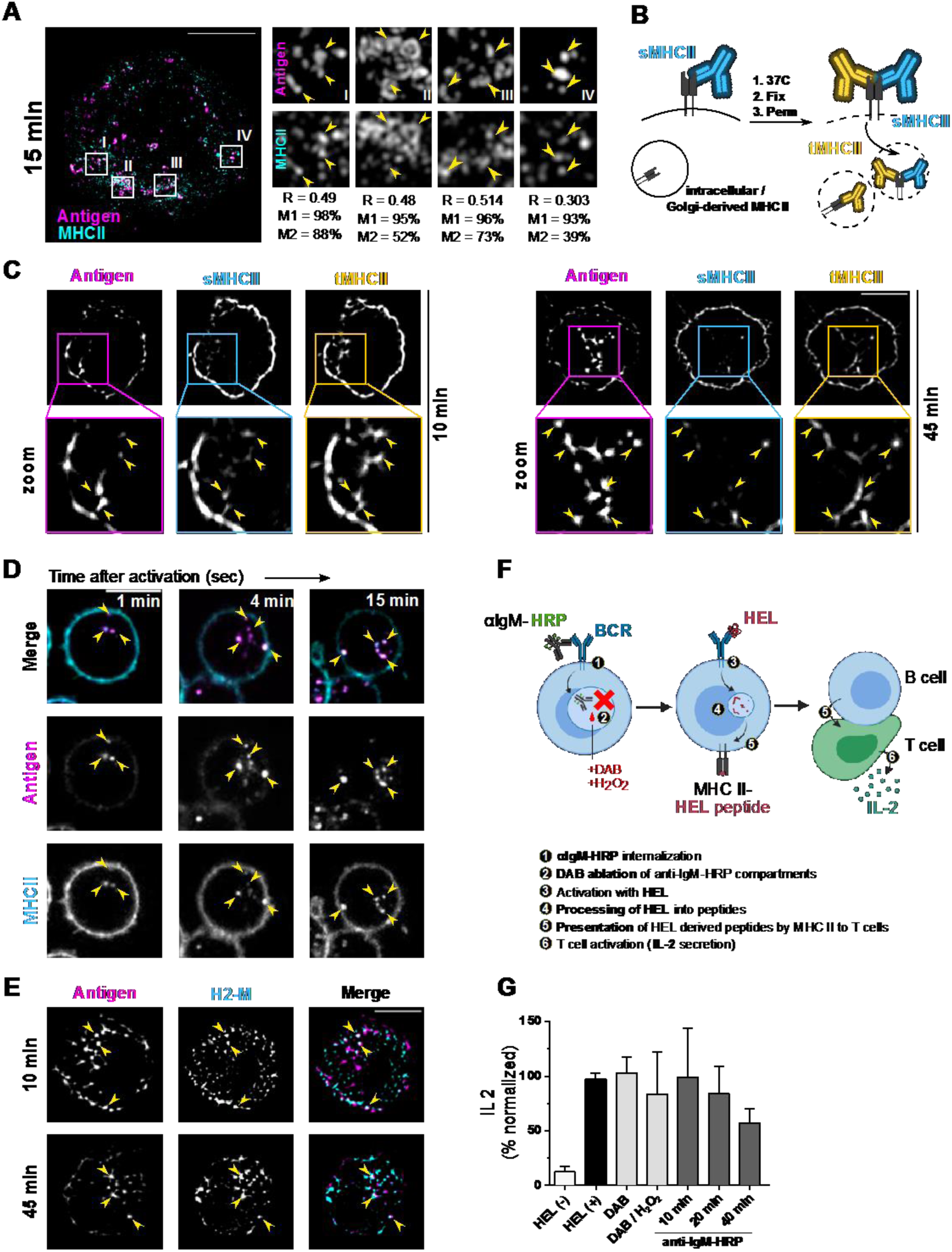
Antigen and surface-derived MHCII rapidly converge after internalisation. **A** SIM imaging of A20 D1.3 cells activated with AF647-αIgM (antigen, magenta) for 15 min and immunostained for MHC-II (cyan). A representative cell (scale bar 5 µm; stack image; 0.125 µm step size) is shown on the left with white squares indicating insets I-IV (1.6 µm x 1.6 µm) shown on panels on the right. Quantification of each inset is shown below as M1 (Manderścoefficient 1; % colocalisation of antigen with MHCII), M2 (Manderścoefficient 2; % colocalisation of MHCII with antigen) and R (Pearsońs correlation coefficient). **B** A schematic view on the staining to distinguish surface-derived MHCII from the total pool, used in C-D. **C** A20 D1.3 cells (plane image; antigen in magenta) were stained with anti-MHCII (AF488) before activation with RRx-αIgM (antigen, magenta) to label surface-bound MHCII (sMHCII, cyan). After activation for 10 or 45 min at 37°C, cells were fixed and permeabilised to stain with anti-MHCII and a secondary antibody (AF633; tMHCII, yellow). Samples were imaged with iterative imaging of a single plane with SDCM (20-25 frames/plane) and post-processed to obtain SRRF super-resolution image. Upper panel: representative cell; lower panel, zoom-in of the white square in the upper panel. Examples of colocalising vesicles are pointed with yellow arrowheads. Scale bar 5um. **D** Live imaging of A20D1.3 stained on ice with AF488-anti-MHCII (cyan) and RRx-αIgM (antigen, magenta). Samples were imaged every 5 seconds using SDCM after 1 min at 37 °C (ORCA camera). A timelapse from a representative cell is shown and examples of colocalising vesicles are pointed with yellow arrowheads. Scale bar 5 µm. See Movie S4. **E** SRRF imaging of A20 D1.3 cells activated with AF647-αIgM (antigen, magenta) for 10 or 45 min and immunostained for H2-M (cyan). A representative cell is shown and examples of colocalising vesicles are pointed to with yellow arrow-heads. Scale bar 5 µm. **F** A Schematic illustration explaining the experimental process of DAB-mediated endosome ablation and antigen presentation to T cells. **G** Effect of DAB mediated endosome ablation on antigen presentation measured as IL-2 secretion by ELISA, as schematically illustrated in F. HEL (-): negative control, untreated A20 D1.3 cells without antigen. HEL (+): positive control, untreated cells activated with HEL. DAB: cells treated with DAB and HRP (without H2O2) and activated with HEL. DAB/H2O2: cells treated with DAB and H2O2 (without HRP) and activated with HEL. HRP-αIgM: cells activated with HRP-αIgM for different time points, treated with DAB and H2O2 and activated with HEL. Results (mean ± SD, n = 3) are shown as % of IL-2 secretion normalised to the control cells (HEL(+), 100%).

**Figure 6.**
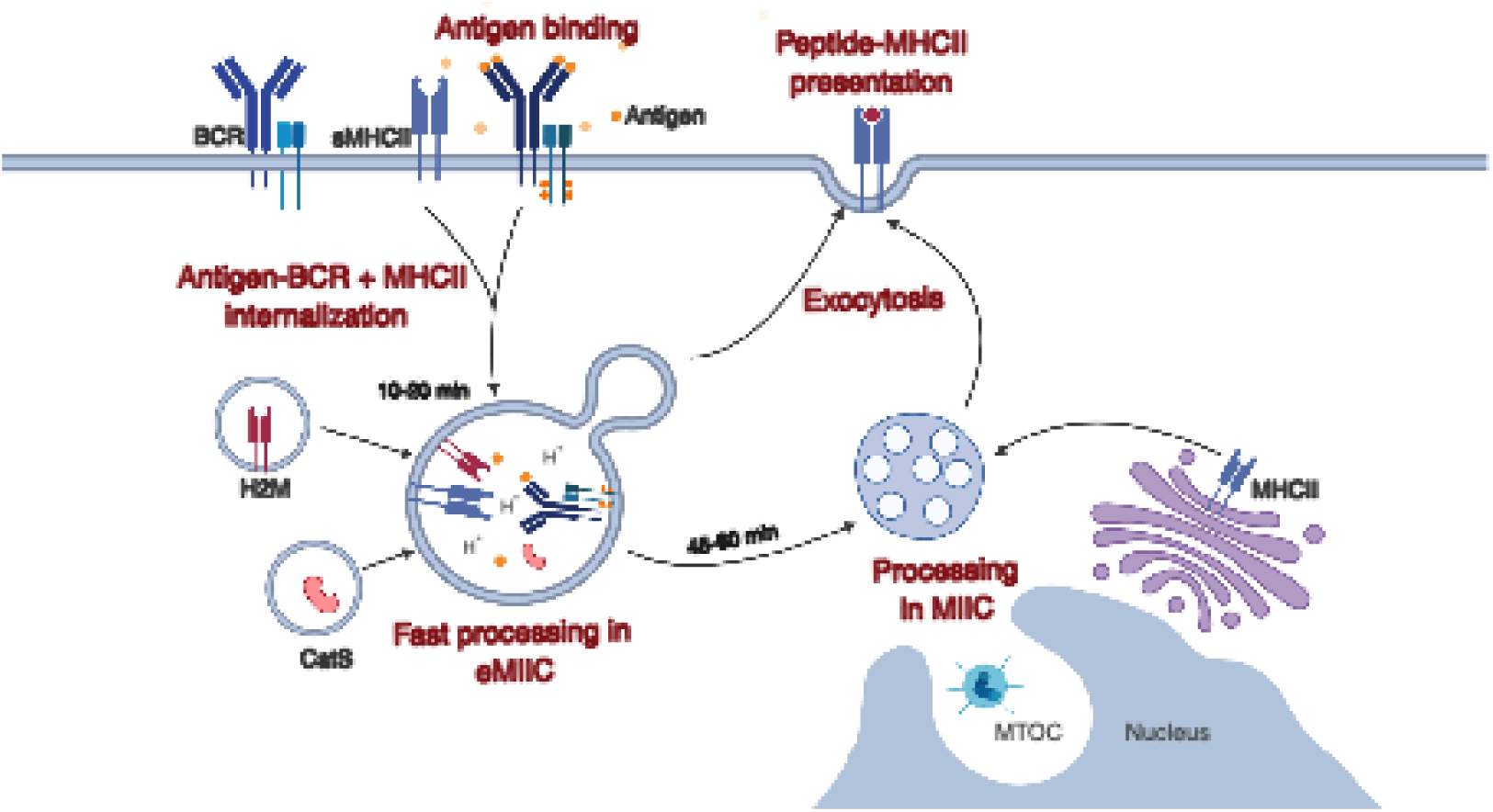
Model of antigen processing in B cells. B cell internalise antigen and surface MHCII (sMHCII) and target them to early MHCII Compartments (eMIIC) to support fast antigen processing and pMHCII presentation. At later stages, antigen is targeted to classical MIIC compartments in the perinuclear region for further pMHCII presentation.

Finally, to examine the functionality of eMHCII compartments in peptide loading, we utilized a well-stablished 3,3’ diaminobenzidine-peroxidase (DAB-HRP) endosome ablation technique combined with an ELISA-based antigen presentation assay. Monomeric DAB is polymerised in the presence of H_2_O_2_ and HRP, selectively fixing the HRP-containing endosomes by crosslinking the luminal and membrane integral proteins of the vesicles (Henry and Sheff, 2008; Pond and Watts, 1999; Stoorvogel et al., 1996). We pulsed A20 D1.3 cells with HRP-labeled αIgM (HRP-αIgM) for 10, 20 or 45 minutes followed by DAB-ablation of the HRP-αIgM-containing endosomal compartments. Then, we activated the endosome-ablated A20 D1.3 cells with HEL antigen. Using a cognate T cell line 1E5, that recognizes HEL-derived peptides on *I-Ad* MHCII of the A20 D1.3 B cells, we were able to measure IL-2 secretion by T cells as an antigen presentation readout. We found normal levels of peptide presentation when HRP-αIgM vesicles were ablated after 10 minutes, and close to normal levels after 20 min activation. In these time points, however, only a fraction of the antigen is internalized (Fig. S2D), and the cells continue internalizing antigen and are likely to have remaining endosomal capacity for trafficking. This indicates that, even if all HRP-αIgM endosomes are ablated at these time points, a pool of early carriers still remains functional. Interestingly, we also detected robust presentation, 60% compared to the non-treated cells, in cells where HRP-αIgM vesicles were ablated after the 40 minutes. In this time point, most of the antigen has been internalized and reached the perinuclear MIICs (see Fig. 1 and Fig. S2D). This result suggests that the perinuclear MIICs are not fundamentally required for presentation but the eMIIC could also generate pMHCII to support pMHCII generation. Nevertheless, it is also possible that the DAB-HRP reaction does not completely abolish all perinuclear MIICs, or that they are regenerated during the second activation maturing from a non-ablated endosomal pool.

## Discussion

To the study vesicular networks responsible for antigen processing in B cells, we utilized high and super-resolution microscopy for systematic colocalisation analysis of antigen with key markers of various endolysosomal compartments and known components of MIIC. Consistent with previous studies (Aluvihare et al., 1997; Siemasko et al., 1998; Vascotto et al., 2007a), we observed that, over time, antigen concentrates in the perinuclear region together with LE/MBV markers LAMP1, Rab7 and Rab9, in compartments well-fitting to the description of MIIC. However, we also observed fast and highly efficient targeting of antigen into acidic compartments, that also possessed key features of MIIC, already in minutes after internalisation. These vesicles, located in the cell periphery, displayed a heterogenous combination of early and late endosomal markers and also exhibited variable ultrastructural morphologies. Interestingly, we show robust recruitment of surface-derived MHCII to these compartments, that we named eMIICs, suggesting that they could support fast presentation using MHCII recycled from the plasma membrane. This work provides the first endosomal roadmap of the intracellular trafficking of antigen in B cells and reveals previously unappreciated efficacy in MIIC formation.

Much of our knowledge in B cell antigen processing compartments is derived from biochemical studies, including cell fractionations, radiolabelling of antigen, and electron microscopy (Amigorena et al., 1994; Lankar et al., 2002; West et al., 1994). While already these early studies drew a valid picture of late endosomal or lysosomal, i.e. LAMP1-positive, multivesicular compartment, the approaches were not suitable to address questions about intracellular localization or dynamics of the antigen vesicles. Yet, these features have been strongly linked to distinct functional properties of endolysosomes and they also inform us about the possible molecular machineries regulating the vesicle traffic (Huotari and Helenius, 2011; Hutagalung and Novick, 2011). Our microscopic analysis revealed a remarkable heterogeneity in the endolysosomal markers of antigen vesicles (Fig. 1-3). However, overlapping fluorescent signals could be derived from a vesicle containing two markers, two vesicles containing different markers, or a multilobular vesicle with distinct markers in different domains, not resolvable by conventional light microscopy. Therefore, the small size and crowdedness of the vesicles generated challenges for the colocalisation analyses, particularly affecting Manderś overlap coefficient, which relies on area overlap. We could, at least partially, overcome by the super-resolution SRRF and SIM analyses. While SDCM can achieve a lateral resolution of 250-300 nm, SIM and SRRF improve the x-y resolution by approximately 2-fold. SIM also improves the axial resolution by 2-fold from approximately 600 to 300nm.

The vesicle heterogeneity could be linked to the notion that antigen enters vesicles with low pH (indicated by LysoTracker and fast decay of FITC fluorescence), and degradative capacity (demonstrated by DQ-Ova signal and partial overlap with CatS) extremely fast after internalisation (Fig. 4). It has also been shown that the amounts of Rab-proteins on a given vesicle can fluctuate, increasing the noise in the colocalisation parameters (Huotari and Helenius, 2011; Hutagalung and Novick, 2011; Rink et al., 2005; Vonderheit and Helenius, 2005). Notably, we found that antigen also trafficked in atypical vesicles stably marked by both early endosomal Rab5 and LysoTracker indicating that the heterogeneity of the vesicles would be a more constant feature and not a mere transition state. Our data does not clearly fit the classical “Rab conversion” model, where a vesicle rapidly shifts from Rab5-positive into Rab7-positive (Huotari and Helenius, 2011; Hutagalung and Novick, 2011). Instead, the data might better comply with an alternative model, where sequential budding of membrane domains with LE markers would occur from EE/sorting endosomes (Huotari and Helenius, 2011; Wandinger-Ness and Zerial, 2014) and, indeed, we often detected adjacent localization of different markers possibly indicative of distinct domain on the same vesicle.

Martinez-Martin and colleagues used SIM to demonstrate, in primary B cells, that 15 min after activation, part of the internalised antigen concentrated in ring-like structures representing autophagosomes (Martinez-Martin et al., 2017). However, it remains unclear what could be the role of autophagy in terms of antigen fate or pMHCII processing. In our SIM analysis, we also detected some ring-like structures, that could represent autophagosomes (Fig. 5A) and the partial partitioning of antigen in these autophagosomes, or amphisomes, could explain some of the vesicle heterogeneity we observed. Our data also does not rule out contribution of other vesicular carriers, like clathrin-independent carriers (CLICs), void of specific markers (Kirkham et al., 2005).

An interesting finding from our live imaging data was that the LysoTracker positive, i.e. low pH vesicles, appeared to hover close to the plasma membrane and capture antigen right after internalisation (Fig. 4F). Some of these LysoTracker-positive vesicles also contained Rab5 already before cell activation (Fig. S6C). The overexpression of Rab-proteins has some caveats and Rab5-GFP can, for instance, generate enlarged EEs and lead into only partial recapitulation of endogenous Rab5. We, however, also stained B cells for different pairs of endogenous early and late endosomal markers, and consistently found indications of colocalisation of both markers, especially in early stages after cell activation but to some extent already prior to cell activation (Fig 3F; Fig. S6). This effectiveness suggests prewiring of the B cells endolysosomal system towards antigen presentation, accompanied or boosted by a signalling component from the BCR, as indicated by our analysis in steady state vs activated cells (Fig. S6) and suggested already by Siemasko and colleagues (Siemasko et al., 1998). As such, we support MIIC to be considered as a member of the growing family of specialized endolysosome-related organelles (ELRO) with diverse functions, as proposed in a recent review by Delevoye and colleagues (Delevoye et al., 2019). Considering the poor compliance of antigen vesicles with classical endolysosomal pathway, other ELROs could serve as valuable additional points of comparisons for studies of MIIC membrane traffic. It has been shown that B cells on activatory surfaces mimicking immunological synapses, polarize the MTOC and acidic MHCII vesicles to secrete proteases for antigen extraction (Yuseff et al., 2011). While this happens at later stages of activation and is proposed to precede antigen internalisation, it demonstrates atypical functions of B cell acidic compartments, perhaps analogous to the secretion of lytic granules, another type of ELRO, by CD8+ T cells (Delevoye et al., 2019; Yuseff et al., 2013).

Early biochemical studies, using lipopolysaccharide-activated B lymphoblasts, have proposed the existence of peptide-loaded MHCII in multiple endolysosomal compartments (Castellino and Germain, 1995) and, using the same B cell line than us, demonstrated that B cells can indeed present antigen already in 20 min after activation (Aluvihare et al., 1997). Furthermore, studies have shown antigen degradation into peptides 20 min after activation (Barroso et al., 2015; Davidson et al., 1990). These studies are consistent with our finding that the internalised antigen vesicles highly efficiently colocalise with MHCII in various compartments, as well as partially overlap with Cathepsin-S and H2-M (Fig. 4-5). The acidic nature and degradative capacity of the eMIICs (Fig. 4-5) further supports function in antigen processing, which is also suggested by robust pMHCII presentation detected in the cells where late perinuclear MIIC were ablated with DAB-HRP reaction (Fig 5F).

Interestingly, we found that the newly internalised antigen robustly co-localized with surface-derived MHCII (Fig. 5), suggesting that the pre-existing pool of MHCII could be used for the first wave of pMHCII presentation. This point has been previously tested using cycloheximide, known to block de novo protein synthesis. There, however, cycloheximide was found to inhibit all presentation and it was interpreted so that B cells could only present peptides on newly synthesized MHCII (Aluvihare et al., 1997). Later, concerns have been raised on the side effects of cycloheximide. These include disturbance of vesicle trafficking, actin cytoskeletal dynamics and cell polarization and motility (Clotworthy and Traynor, 2006; Darvishi and Woldemichael, 2016; Oksvold et al., 2012). Thus, the old findings with cycloheximide warrant for a revisit with a sensitive pulse assay for antigen presentation together with more specific inhibitors like, for example, the newly developed FLI-06 that targets ER-exit sites and trans-Golgi network (Yonemura et al., 2016). Our suggestion that biosynthetic MHCII probably arrives to MIIC at later stages, is supported by the old metabolic labelling studies, where, again using the same B cell line than in our study, it was shown that the newly synthesized MHCII arrives to MIIC in 30-60 min after cell activation (Amigorena et al., 1994).

eMIICs could facilitate the speed of pMHCII presentation but could also tune the peptide repertoire. In cell fractionation studies, an MHC class II-like protein H2-O has been reported to concentrate more with the EE fraction as compared to LE fraction, while the peptide-loading chaperone H2-M shows the opposite trend (Gondré-Lewis et al., 2001). While H2-O has been characterized with an inhibitory effect on H2-M, it has also been shown to modulate the repertoire of peptides presented on MHCII with a mechanism still unclear (Denzin et al., 2005; Karlsson, 2005). Due to lack of working antibodies for mouse cells, we were not able to analyse H2-O in our system. Nevertheless, H2-O has been shown to dissociate from H2-M in acidic pH, thereby releasing the inhibition of peptide loading by H2-M (Jiang et al., 2015). This mechanism would allow peptide loading already from eMHCII even in the presence of H2-O. Different ratios of H2-M and H2-O could thus distinguish the peptides sent out from eMIICs from those originating from mature MIIC.

Using TEM, we found antigen in vesicles with diverse morphologies (Fig. 1F, left; Fig. S1A). Ranging from spherical to multilobular, various compartments harboured intralumenal vesicles, consistent with reports characterizing MIICs with multivesicular features (Roche and Furuta, 2015; Unanue et al., 2016; van Lith et al., 2001; Xiu et al., 2011). Antigen-containing single-membrane vesicles with round or horse-shoe shapes were also detected. While it has been shown in dendritic cells that intralumenal vesicles are not required for MHCII loading (Bosch et al., 2013), also multilamellar MIIC have been reported (Unanue et al., 2016). These notions suggest that MIIC function is not bound to certain vesicle morphology. Based on both the morphological and vesicle marker-based heterogeneity, we propose that early peripheral antigen vesicles, eMIICs, are functional MIIC in transit. While eMIICs might part off and fuse again or migrate as such to the perinuclear region for gradual maturation into MIIC, we suggest that they are functional throughout the pathway.

## Acknowledgements

We are thankful for Laura Grönfors and Mervi Lindman for technical assistance. Microscopy and flow cytometry were performed at Turku Bioscience Cell Imaging and Cytometry (CIC), supported by Turku Bioimaging and Euro-Bioimaging, thanked for generous help and expertise. Biocenter Finland is acknowledged for providing research infrastructures, particularly at CIC and the Electron Microscopy Unit, Institute of Biotechnology, Helsinki. We thank Tampere Imaging Facility for sharing their image analysis resources. Prof. Johanna Ivaska and Dr. Pranshu Sahgal are acknowledged for their help and generosity regarding reagents and protocols, and Prof. Johanna Ivaska and Prof. Ari Helenius for constructive discussions. Juan Palacios-Ortega is acknowledged for help in manuscript formatting and sharing reagents.

## Materials and Methods

For more information about reagents and antibodies, please check Table S1.

### Cells and mice

A20 mouse lymphoma cells stably expressing a hen egg lysozyme (HEL)–specific IgM BCR (D1.3) (Williams et al, 1994) and 1E5 T cells, kind gifts from Prof Facundo Batista, stably expressing a transgenic TCR specific for HEL_108–116_/*I-Ad* (Adorini et al., 1993) were maintained in complete RPMI (cRPMI; RPMI 1640 with 2.05 mM L-glutamine supplemented with 10% fetal calf serum (FCS), 50 μM β-mercaptoethanol, 4 mM L-glutamine, 10 mM HEPES and 100 U/ml Penicillin/Streptomycin). Cells were regularly examined for bacterial and fungal contaminations and tested for mycoplasma contaminations, but no other tests were run on the cell lines. Primary splenic B cells were isolated from 2-5 months-old male and female MD4 mice (C57BL/6-Tg(IghelMD4)4Ccg/J, The Jackson Laboratory) using a negative selection kit (StemCell Technologies, #19854). All animal experiments were approved by the Ethical Committee for Animal Experimentation in Finland. They were done in adherence with the rules and regulations of the Finnish Act on Animal Experimentation (62/2006) and were performed according to the 3R-principle (animal license numbers: 7574/04.10.07/2014, KEK/2018-2504-Mattila, 10727/2018).

### Transfection

A20 D1.3 cells were transfected as previously described (Sustar et al., 2018). Briefly, 2 million cells were resuspended in 180ul of 2S transfection buffer (5 mM KCl, 15 mM MgCl2, 15 mM HEPES, 50 mM Sodium Succinate, 180 mM Na_2_HPO_4_/ NaH_2_PO_4_ pH 7.2) containing 2 µg of plasmid and electroporated using AMAXA electroporation machine (program X-005, Biosystem) in 0.2 cm gap electroporation cuvettes. Cells were then transferred to 2ml of cRPMI to recover overnight. Rab5a-GFP plasmid was a kind gift from Prof. Johanna Ivaska.

### B cell activation and visualization of antigen vesicles by immunofluorescence

A20 D1.3 or isolated primary B cells were activated with 10μg/ml of Alexa Fluor-647 or Rhodamine Red-X (RRx) anti-mouse IgM (α-IgM) (Jackson ImmunoResearch), unless indicated otherwise. Cells were labelled with fluorescently-labelled α-IgM for 10 min on ice, washed with PBS to remove excess unbound antigen and resuspend in Imaging Buffer (PBS, 10% FCS). When indicated, cells were also labelled with anti-MHCII-Alexa Fluor 488 on ice. After washing, cells were activated for different timepoints in an incubator (5% CO_2_, 37°C) in a 12-wells PTFE diagnostic slide (Thermo, #10028210), coated with fibronectin, and fixed with 4% PFA 10min at RT. Samples were blocked and permeabilized with blocking buffer (5% horse or donkey serum, 0.3% Triton X100 in PBS) for 20min at RT. After blocking, samples were stained with primary antibodies for 1h at RT or 4°C O/N in staining buffer (1% BSA, 0.3% Triton X100 in PBS), followed by washes with PBS and incubation with the secondary antibodies 30min at RT in PBS. Samples were mounted using FluoroMount-G containing DAPI (Thermo #00495952).

### Visualization of antigen vesicles by live imaging

A20 D1.3 cells (1 million/ml) were labelled with 125 nM LysoTracker Deep Red (Thermo # L12492) for 1 hour in an incubator (5% CO_2_, 37°C), washed with PBS and resuspended in cRPMI. Cells were then labelled with 10μg/ml of donkey anti-mouse IgM-AF488 on ice for 10 min and washed with cold PBS. For surface-MHCII internalisation experiments, cells were stained on ice with anti-MHCII- AF488 and 10μl/ml donkey-anti-mouse IgM-RRx for 5min and washed with cold PBS. Cells were resuspended in cold Imaging Buffer and seeded on 4-well MatTek dishes on ice. After seeding, cells were activated at 37 °C inside the environmental chamber of the microscope and image immediately.

### Image acquisition and processing, spinning disk confocal microscopy

Images were acquired using a 3i CSU-W1 spinning disk equipped with 405, 488, 561 and 640 nm laser lines and 510-540, 580-654 and 672-712 nm filters and 63x Zeiss Plan-Apochromat objective. Hamamatsu sCMOS Orca Flash4 v2 C11440-22CU (2048 x 2048 pixels, 1×1 binning) was used to image fixed samples unless otherwise indicated, and Photometrics Evolve 10 MHz Back Illuminated EMCCD (512 x 512 pixels, 1×1 binning) camera was used to image live samples.

All SDCM images were deconvolved with Huygens Essential version 16.10 (Scientific Volume Imaging, The Netherlands, http://svi.nl), using the CMLE algorithm, with Signal to Noise Ratio of 20 and 40 iterations. For SRRF, 20-50 images were acquired from one single plane using timelapse mode and processed in Fiji ImageJ using the SRRF module.

### Colocalisation analysis

Colocalisation on Spinning Disk Confocal Microscope images were analysed with Huygens Essential version 16.10 (Scientific Volume Imaging, The Netherlands, http://svi.nl), using optimized, automatic thresholding. Colocalisation on SRRF images was performed on ImageJ using Colocalisation Threshold tool. Graphs and statistics were prepared on GraphPad Prism (GraphPad Software, La Jolla California USA).

### Analysis of antigen clustering

Cluster analysis of the deconvolved data was done by batch processing in MATLAB R2018b (The MathWorks Inc.). Binary masks were created from full volumes containing one cell using the method by Otsu. Objects were then segmented in 3D using the regionprops function. Only objects inside a circular mask were kept in order to exclude clusters from adjacent cells, for simplicity this was done in 2D by manually overlaying the image with a circle. The MTOC channel was segmented in the same way and the cluster with the highest intensity value was identified as MTOC. The distances of each cluster to the MTOC was calculated from the centroid positions in 3D. Graphs and statistics were prepared on GraphPad Prism. The scripts can be found on MattilaLab’s GitHub (https://github.com/mattilalab/hernandez-perez-et-al-2019).

### Structured illumination microscopy (SIM)

The samples were prepared as above in “*B cell activation and visualization of antigen vesicles by immunofluorescence”* on fibronectin-coated MatTek dishes and mounted in Vectashield (Vector Laboratories, US) mounting medium. 3D structured illumination (SIM) Imaging was performed with GE Healthcare, DeltaVision OMX SR V4 with 60x/1.42 SIM Olympus Plan Apo N objective, front Illuminated sCMOS cameras, 488, 568 and 640 nm solid-state lasers by optical sectioning of 0.125 µm. The SIM reconstruction was performed with OMX Acquisition software version 3.70. (GE Healthcare, UK).

### Antigen internalisation for flow cytometry

A20 D1.3 cells were stained on ice for 10 min with anti-IgM-biotin (Southern Biotech) or HEL-biotin and washed with PBS. Cells were incubated at 37C and 5% CO2 at different timepoints. For time 0 the samples were kept on ice all the time After incubation, cells were kept on ice and stained with streptavidin-633 (LifeTechonologies #S-21375) for 20min, washed and analysed. BD LSR Fortessa analyser equipped with four lasers (405, 488, 561 and 640nm) was used. Data was analysed using FlowJo v10 (Tree Star).

### Antigen internalisation, immunofluorescence

A20 D1.3 cells were stained on ice for 10 min with biotinylated anti-IgM-Alexa Fluor F647- (labelled in-house) and washed with PBS. Cells were resuspended in Imaging Buffer (PBS, 10% FCS) and activated for different timepoints in an incubator (5% CO2, 37°C) on fibronectin-coated 12-well microscope slide. After activation, slides were kept on ice to stop internalisation and stained with streptavidin-Alexa Fluor 488 (#S11223) for 10 min. Cells were washed with PBS and fixed with 4% PFA 10 min at RT. Samples were mounted using FluoroMount-G (Thermo 00-4958-02).

### DQ-Ova proteolysis reporter

DQ Ovalbumin (Thermo Fisher Scientific D12053) was biotinylated in-house with EZ-Link Maleimide-PEG2-biotin (Thermo 21901BID). HEL from (#L6876 Sigma) was biotinylated using EZ-Link™ Sulfo-NHS-LC-LC-Biotin (Thermo 21338). A20 D1.3 cells were first incubated with 10 μg/ml biotin-HEL or biotinylated anti-IgM (Southern Biotech) for 10 min on ice. After washing with PBS, cells were incubated for 5min on ice with unlabelled streptavidin for IF samples or Alexa Fluor 633-labelled streptavidin for flow cytometry samples, wash with PBS, and incubated 5min on ice with biotinylated-DQ-Ova. After 3 washes with PBS, cells were activated in an incubator (5% CO2, 37°C) to allow internalisation of the probe-linked antigen. After the activation, cells were placed on ice and analysed by flow cytometry immediately. For immunofluorescence samples, cells were activated on 12-well slides coated with fibronectin in the incubator, fixed with 4% PFA after activation, and stained as previously described. DQ-Ova was excited with 488 nm laser and measured with filters identical to Alexa Fluor 488 or GFP.

### Assessment of low pH for flow cytometry

A20 D1.3 cells were stained on ice for 10 min with anti-IgM-AF647 (5 ug/ml) and anti-IgM-FITC (5 ug/ml) and washed with PBS. Cells were then incubated at 37C and 5% CO2 at different timepoints in a 96 well-plate for flow cytometry analysis. After incubation, cells were kept on ice and analysed using a BD LSR Fortessa analyser equipped with four lasers (405, 488, 561 and 640nm). As time 0, the samples were kept on ice all the time. Data was analysed using FlowJo v10 (Tree Star).

### Transmission electron microscopy

A20 D1.3 cells were activated with a mixture of 6 nm colloidal-gold conjugated goat anti-Mouse IgM (Jackson ImmunoResearch, 115-195-075; 1:650 dilution) and 20 μg/ml Alexa Fluor 647 labelled donkey anti-mouse IgM F(ab’)_2_ fragments (Jackson ImmunoResearch, 715-606-020) in imaging buffer (0.5mM CaCl_2_, 0.2mM MgCl_2_, 5.5mM D-Glucose, 10% FBS in PBS) and placed on fibronectin (4 μg/ml) coated glass coverslips (thickness #1) for 15 or 75 min. The cells were fixed with 2 % Glutaraldehyde (EM-grade, Sigma G7651) in 0.1 M Na-Cacodylate buffer, pH 7.4, for 30 min at room temperature, and then washed twice for 3 min with 0.1 M Na-Cacodylate buffer, pH 7.4. The samples were processed for TEM as described in Seemann et al., 2000. 60-nm-thick sections parallel to the cover slip were cut using a Leica EM Ultracut UC7 ultramicrome (Leica Mikrosysteme GmbH, Austria). The electron micrographs post-stained with uranyl acetate and lead citrate, and imaged with Jeol JEM 1400 transmission electron microscope (Jeol Ltd., Tokyo, Japan) equipped with a bottom mounted CCD-camera (Orius SC 1000B, Gatan Inc., Pleasanton, CA) and Jeol JEM- 1400 Plus equipped with OSIS Quemesa bottom-mounted CCD-Camera (EMSIS, Germany), both operating at 80 kV.

### DAB endosome ablation

Endosome ablation assay was adapted from Pond and Watts, 1999. A20 D1.3 cells (10_7_/ml) were incubated in FCS-free RPMI for 45 minutes at 37°C. Cells were surface-stained on ice with 10 μg/ml of anti-IgM-biotin (Southern Biotech) for 10 minutes, followed by one PBS wash. Then, cells were incubated with streptavidin-HRP for 10 minutes on ice and washed twice with PBS. Internalization of anti-IgM-HRP was initiated by incubation at 37°C 5% CO_2_ for different times (10, 20 and 40 minutes). After that, vesicle traffic was stopped by incubation on ice and anti-IgM-HRP-containing endosomes were ablated by addition of 0.1 mg/ml DAB (Santa Cruz sc-24982) and 0.025% H_2_O_2_ in freshly prepared DAB buffer (70 mM NaCl, 20 mM HEPES, 2 mM CaCl2 and 50 mM ascorbic acid) for 30 minutes on ice in the dark. Ascorbic acid is a membrane-impermeable molecule that acts as a radical scavenger inhibiting extracellular HRP activity to avoid DAB deposits on the plasma membrane. As a control, cells were incubated in DAB buffer with 0.1 mg/ml DAB, but without HRP or without H_2_O_2_. Cells were then washed 3 times with PBS-1% BSA and kept on ice. Viability after endosome ablation assessed with Trypan Blue was 95-98%.

### Antigen presentation measured by ELISA

After endosome ablation, A20 D1.3 cells were incubated with 10 μg/ml of HEL for 1h at 37°C in cRPMI. After 1h, cells were washed and resuspend in cRPMI. A20 D1.3 B cells were mixed with 1E5 T cells (ratio 2:1) and incubated at 37°C 5% CO2 overnight. IL-2 secretion levels were measured next day by ELISA on half-area 96-well plates coated with capture antibodies (anti-IL-2) for 1h at 37°C in 25 µL of PBS. Non-specific binding sites were blocked overnight at 4°C in 150 µL of blocking buffer (PBS, 1% BSA). Appropriate dilutions of 50 µL supernatant samples in cRPMI were added to the ELISA plate for 1-2h incubation at 37°C. Biotin-conjugated detection antibodies (anti-IL-2-biotin) in 50 µL of blocking buffer are added for 1 hour at RT followed by 50 µL AP (alkaline phosphatase)-streptavidin in blocking buffer for 1 hour at RT. In between all incubation steps, plates were washed with 150 µL washing buffer (PBS, 0.05% Tween-20). The final wash was completed with 2 times wash with 150 µL of water. Finally, 50 µL of pNPP solution was added and optical density (OD) was measured at 405 nm. Typical time for AP-substrate incubation before measurement was about 10-20 min at RT, before reaching signal saturation.

All ELISA samples were run in duplicates, OD values were averaged and blank background was subtracted. All samples were normalized to the control cells (100%) and data is presented as mean values and standard deviation of 3 experiments.

### Statistical analysis and illustrations

Statistical significances were calculated using unpaired Student’s *t*-test assuming normal distribution of the data. Statistical values are denoted as: \**P*<0.05, \*\**P*<0.01, \*\*\**P*<0.001, \*\*\*\**P*<0.0001. Graphs were created in GraphPad Prism 6 and illustrations were created with BioRender. Figure formatting was done on Inkscape v.092.2.

## SUPPLEMENTARY INFORMATION

**Figure S1.**
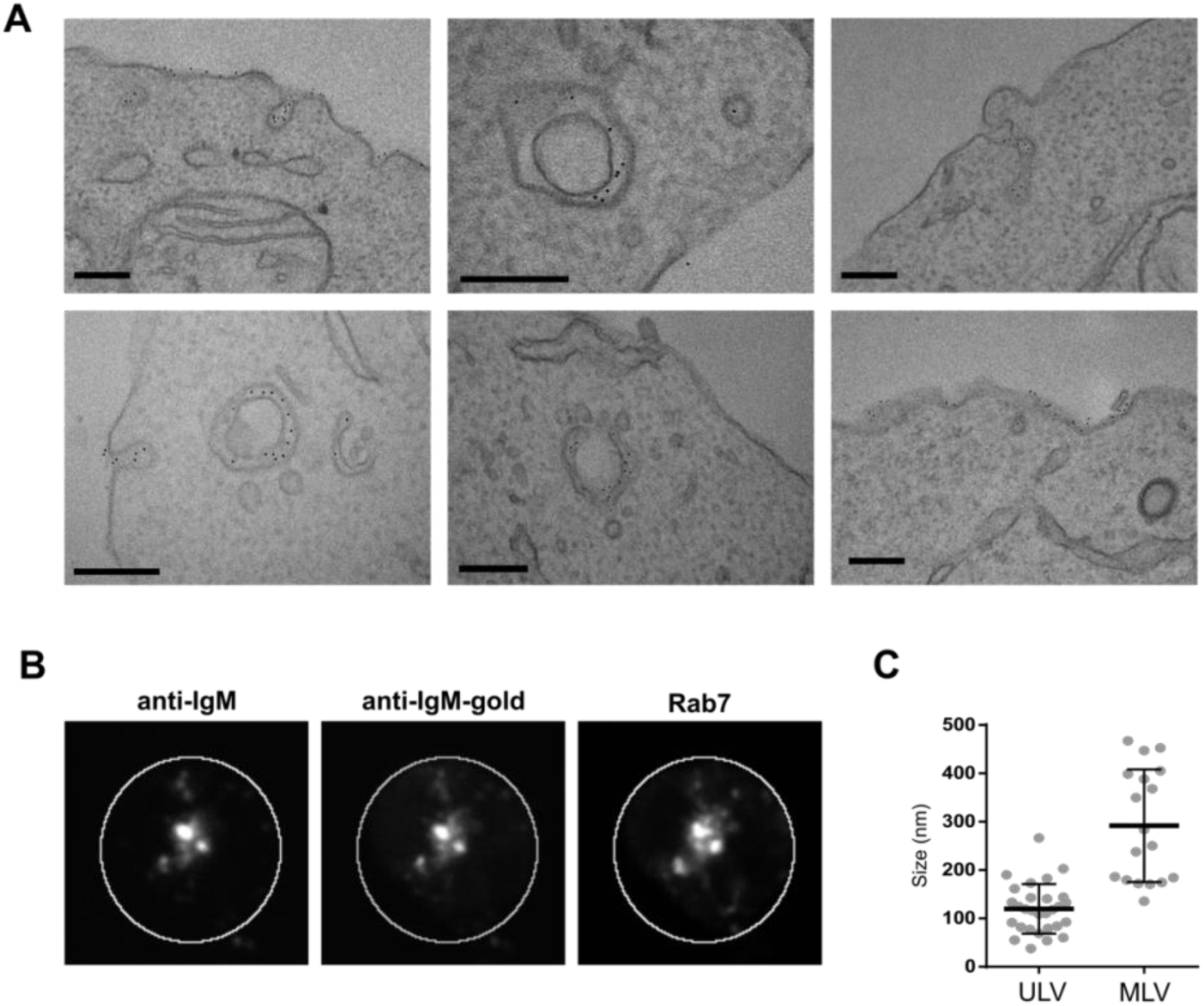
**A** A20 D1.3 cells were activated with αIgM conjugated with 6nm colloidal gold particles mixed with AF647-αIgM, for 15 min and imaged using Transmission Electron Microscopy. Scale bars 200 nm. **B** The possible effect of colloidal gold-conjugation to the localization of α-IgM was controlled by immunofluorescence analysis of sample duplicates (from A). The cells activated for 75 min were fixed and permeabilised, and gold-conjugated αIgM was stained using an isotype-specific secondary antibody (middle panel) and compared to AF647-αIgM (left). The cells were also immunostained with anti-Rab7 antibody (right). SDCM image shows single section of a representative cell (cell plasma membrane marked with a white circle), where a strong colocalisation of fluorescently labelled α-IgM and gold-conjugated α-IgM was detected together with Rab7, similarly to the samples without colloidal gold (Fig. S1B; see Fig. 1A and Fig. 2B). **C** EM micrographs (as in panel A) were subjected to vesicle size analysis. Vesicles were classified as unilamellar (ULV) or multilamellar (MLV) and their diameter was measured in ImageJ.

**Figure S2.**
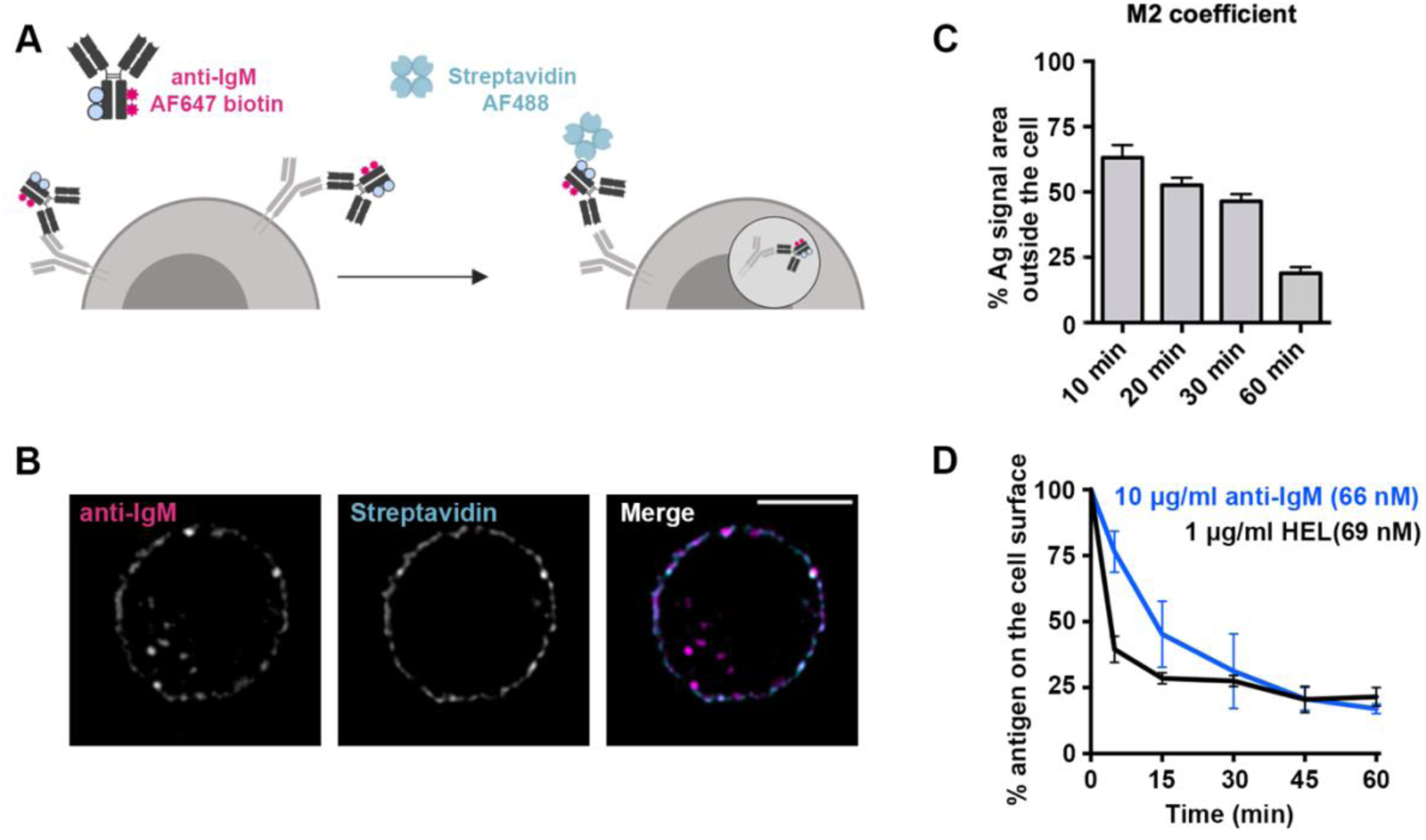
**A** Schematic representation of the staining to distinguish between internalised AF647-αIgM (magenta) and surface-resident αIgM (probed with AF488-streptavidin; cyan). **B** SDCM imaging of A20 D1.3 cells activated with biotin-AF647-αIgM for 10 min. AF647-αIgM used for activation is shown in magenta, and surface-resident αIgM (AF488-streptavidin) in cyan. SDCM images were deconvolved with Huygens software. Single confocal plane from a representative cell is presented. Scale bar 5 µm. **C** Quantification of the data in B, including additional timepoints. 3D images from cells activated for 10, 20, 30 and 60 minutes were analysed for Manderś overlap coefficients (M2) using ImageJ. Data shown as mean ±SEM of one experiment (n>20 cells per timepoint). **D** A20 D1.3 cells labelled with biotinylated-αIgM or biotinylated-HEL were incubated at 37 °C at different timepoints and stained on ice with AF488-streptavidin to detect surface-resident αIgM. Intensity was normalised to time 0 (100%). Data from at least three independent experiments is presented as mean ±SD.

**Figure S3.**
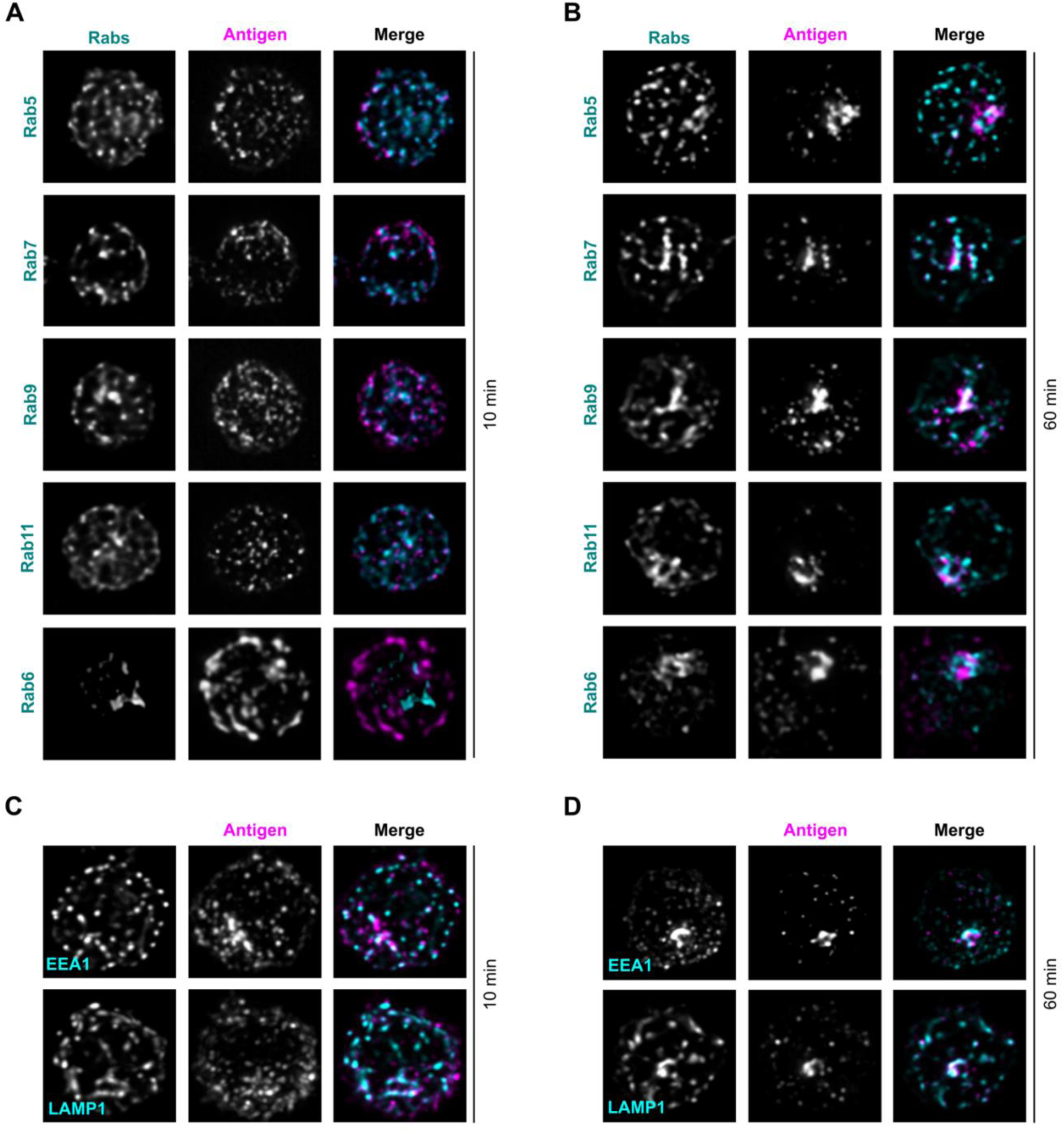
Colocalization of antigen with different Rab-proteins. **A-B** SDCM imaging of A20 D1.3 cells activated with AF647-αIgM (antigen, magenta) for 10 min **(A)** or 60 min **(B)** and immunostained for different Rab-proteins: Rab5, Rab7, Rab9, Rab11 and Rab6 (cyan). Images were deconvolved with Huygens software. Z-projections of the 3D images from representative cells are shown. **C-D** SDCM imaging of A20 D1.3 cells activated with AF647-αIgM (antigen, magenta) for 10 min **(C)** or 60 min **(D)** and immunostained for EEA1 or LAMP1 (cyan). Images were deconvolved with Huygens software. Z-projections of the 3D images from representative cells are shown.

**Figure S4.**
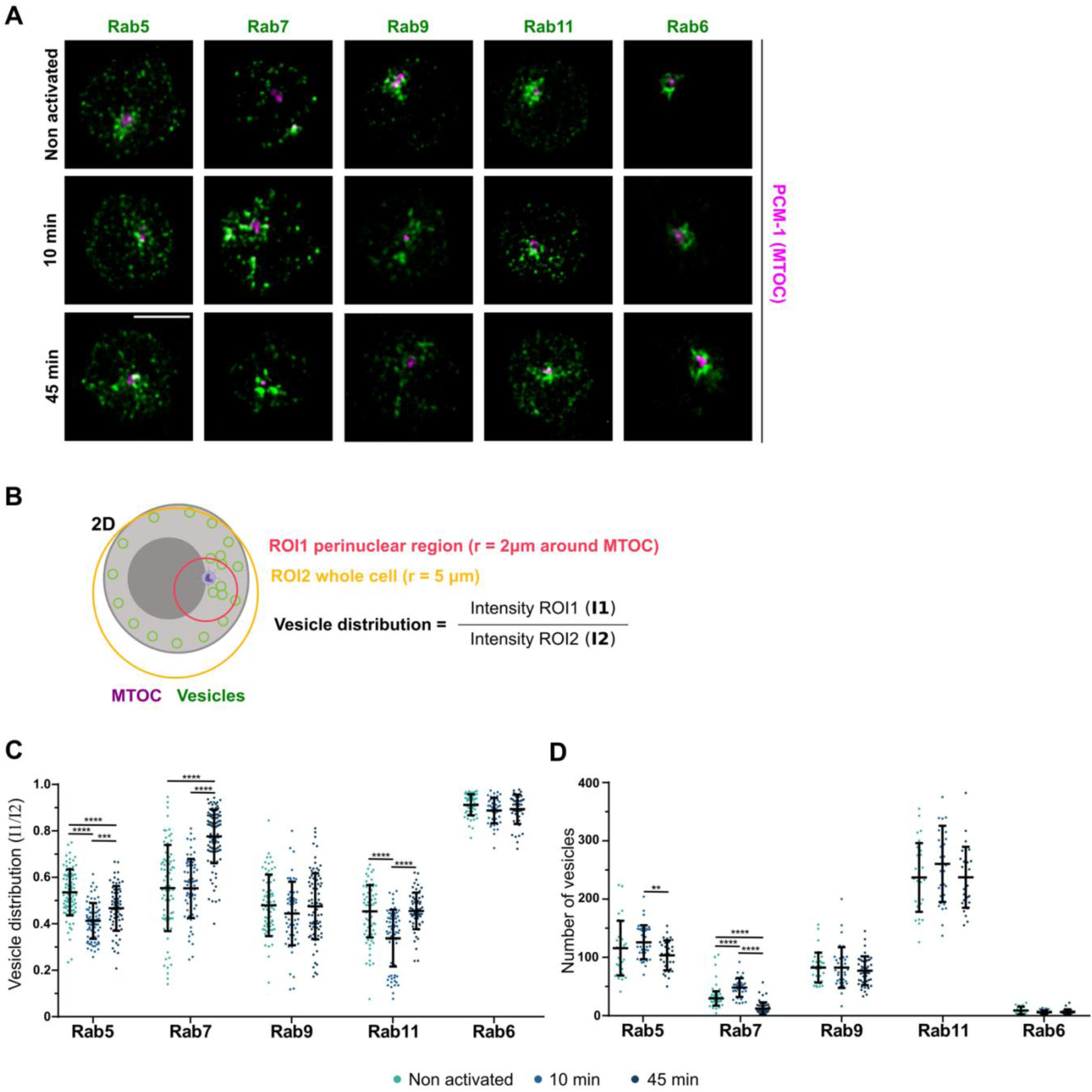
Effect of B cell activation on the distribution of Rab compartments. **A** SDCM imaging of A20 D1.3 cells non-activated or activated with αIgM for 10 min or 45 min and immunostained for different Rab-proteins (Rab5, Rab7, Rab9, Rab11 and Rab6) in green, and an MTOC marker PCM—1 in magenta. Images were deconvolved with Huygens software. Z- projections of the 3D images from representative cells are shown. **B** Schematic representation showing the analyses performed on the images in **A** to generate quantification in **C**. In ImageJ, two different regions of interested (ROI) were selected: ROI1, a circle with radius of 2 μm around the MTOC; and ROI2, a circle around the whole cell. Distribution of the vesicles was quantified as intensity of ROI1/intensity ROI2. **C** Results of the analysis performed as described in B. Data from two independent experiments (mean + SD, 50-100 cells). **D** Results of the quantification of the same data (A) for number of vesicles using the MATLAB script described in Fig.1. Data from one experiment (mean + SD of at least 30 cells).

**Figure S5.**
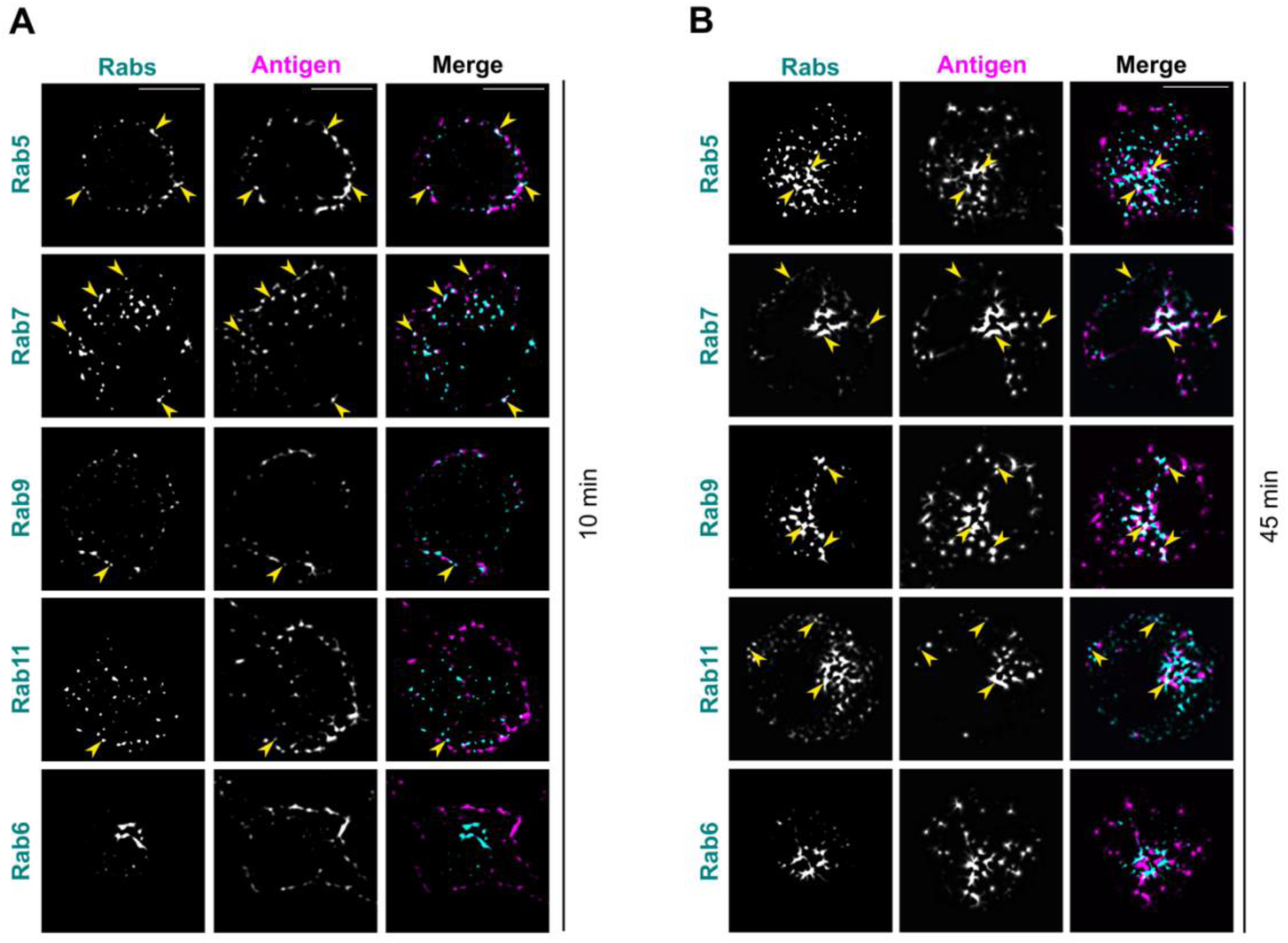
Colocalization of antigen with different Rab-proteins in SRFF. **A-B** SRFF imaging of A20 D1.3 cells activated with AF647-αIgM (antigen, magenta) for 10 min **(A)** or 60 min **(B)** and immunostained for different Rab-proteins: Rab5, Rab7, Rab9, Rab11 and Rab6 (cyan). Z-projections of the 3D images from representative cells are shown. Scale bar: 5 μm.

**Figure S6.**
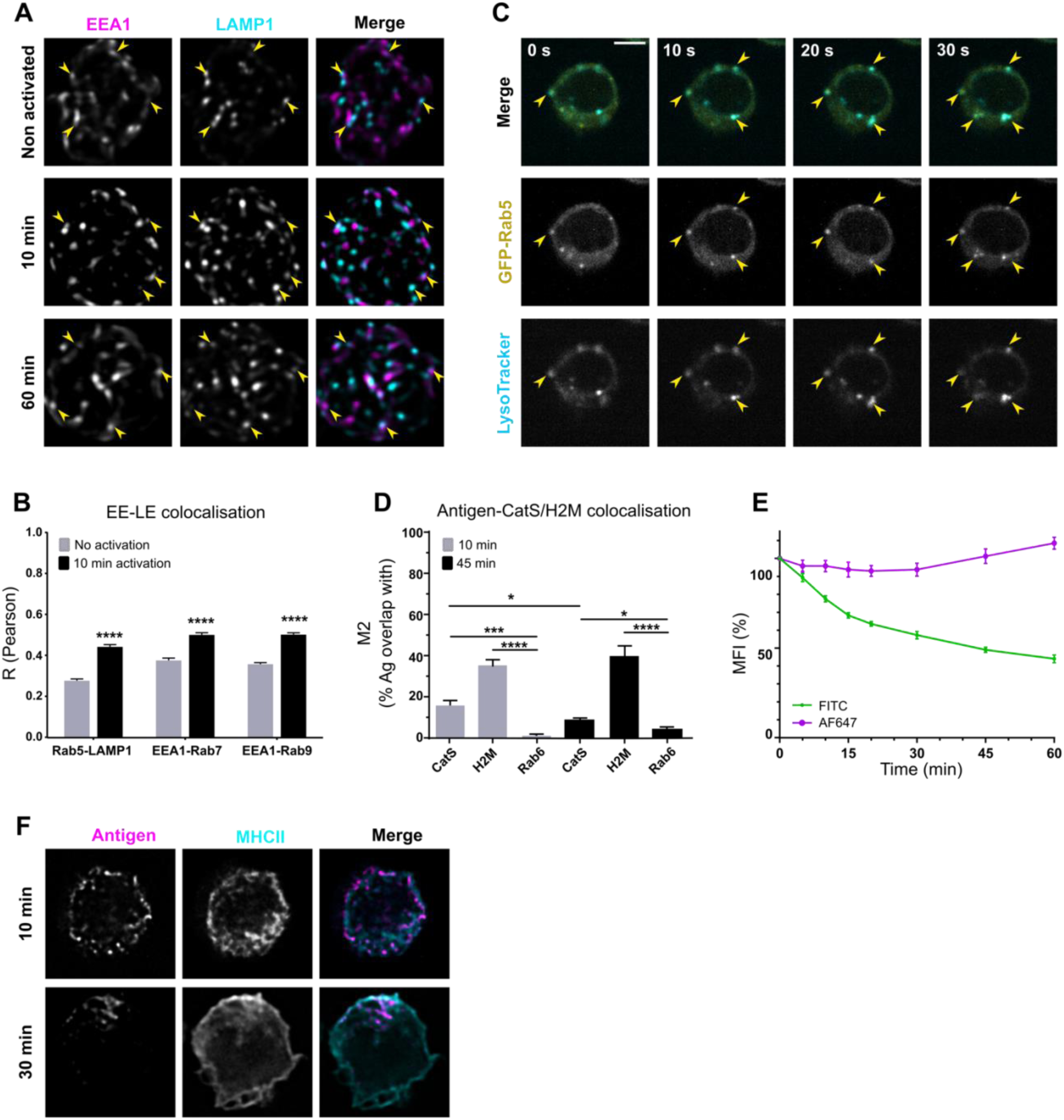
Early and late endosomal markers’ colocalization before and after activation. **A** SDCM imaging of A20 D1.3 cells non-activated or activated with αIgM for 10 min or 60 min and immunostained for EEA1 (magenta) and LAMP1 (cyan). Images were deconvolved with Huygens software. Z-projections of the 3D images from representative cells are shown. **B** Analysis of correlation shown by Pearsońs coefficient of early-late endosomal markers in pairs (Rab5/LAMP1, EEA1/Rab7, EEA1/Rab9) before and after 10 minutes activation. Results from one experiment (n>40 cells) shown as mean +SEM. **C** Rab5-GFP localises to LysoTracker-positive vesicles already before activation. A20 D1.3 cells were transfected with GFP-Rab5 (yellow) and loaded with LysoTracker (LT; cyan). Live-imaging was performed with SDCM (with sCMOS Orca Flash4 v2 camera) on a single plane. On the upper panel, a merge image of a representative cell is shown as a timelapse for 30 seconds. Split channels for GFP-Rab5 and LysoTracker are shown in the middle and bottom panel respectively. Examples of colocalizing vesicles pointed to with yellow arrow-heads. Scale bar 5 µm. **D** Quantification of the data shown in Fig. 4D and Fig. 5E. Antigen colocalization with CatS and H2M, compared to the negative control Rab6, was measured from SRRF images by analysing Manderś overlap coefficients using ImageJ. Data from two independent experiments (>30 cells/timepoint). Results are shown as mean ±SEM. **E** Antigen enters low pH compartments after internalisation. Flow cytometric analysis of pH-sensitive FITC- and pH-stable AF647-conjugated anti-IgM for different timepoints. 3 experiments mean + SD. **F** A20 D1.3 cells were activated with AF647-αIgM (antigen in magenta) for 10 or 30 minutes. Samples were then fixed, permeabilised and stained with anti-MHCII (cyan). SDCM images were deconvolved with Huygens software. Single confocal sections from representative cells are shown.

**Movie S1. Antigen is transported in vesicles positive for early and late endosomal markers.** A20 D1.3 cells were transfected with GFP-Rab5 (right panel), loaded with LysoTracker (left panel) and activated with RRx-αIgM (middle panel). Live-imaging was performed with SDCM (sCMOS Orca Flash4 v2 camera) on a single plane. Triple-positive vesicles are highlighted with a purple circle tracking the spots in the antigen channel. Movie was recorded 10 min after activation and imaged every 2 s.

**Movie S2. Antigen colocalises with acidic vesicles soon after internalisation.** A20 D1.3 cells were loaded with LysoTracker (middle panel) and activated with AF488-αIgM (left panel). Right panel shows merge image of antigen channel (magenta) and LysoTracker channel (cyan). Live-imaging was performed with SDCM (EVOLVE camera) on a single plane every 500 ms. Double-positive vesicles are highlighted with a purple circle tracking the spots in the antigen channel. Movie was recorded 1 min after cell activation.

**Movie S3. Antigen vesicles fuse with LysoTracker-positive vesicles hovering beneath the plasma membrane.** A20 D1.3 cells were loaded with LysoTracker (middle panel) and activated with AF488-αIgM (left panel). Right panel shows merge image of antigen channel (magenta) and LysoTracker channel (cyan). Live-imaging was performed with SDCM (EVOLVE camera) on a single plane every 500 ms. An antigen vesicle fusing with LysoTracker after pinching from the plasma membrane is highlighted with a purple circle. Movie was recorded 80 s after activation.

**Movie S4. Surface MHCII is internalised together with the antigen after activation.** A20 D1.3 cells were labelled on ice with AF488-anti-MHCII (left panel) and RRx-αIgM (middle panel). Cells were shifted at 37 °C to start activation and recorded after 40 s. Right panel shows merge image of antigen channel (magenta) and MHCII channel (cyan). Live-imaging was performed with SDCM (sCMOS Orca Flash4 v2 camera) on a single plane every 4 s. Vesicles positive for MHCII and antigen were tracked with circles (shown in different colours) in the antigen channel.

**Table S1.** Key resources/reagents table

|  | Reagents | Source/Brand | Cat. number | Dilution or concentration | Use |
| --- | --- | --- | --- | --- | --- |
| Antigens | Anti-IgM-biotin | SouthernBiotech | 1021-08 | 10 µg/ml | Antigen internalisation (FACS)<br>DAB ablation |
|  | 6nm Gold rat anti-mouse IgM | Jackson ImmunoResearch (JIR) | 115-195-075 | 1:650 | EM |
|  | Rhodamine Red-X-AffiniPure Donkey anti-mouse IgM | JIR | 715-295-140 | 10 µg/ml | IF/Live imaging |
|  | Alexa Fluor 647AffiniPure Donkey anti-mouse IgM | JIR | 715-605-140 | 5-10 µg/ml | IF/Live imaging/FACS |
|  | Alexa Fluor 488 AffiniPure F(ab')<sub>2</sub> Fragment Donkey anti-mouse IgM | JIR | 715-546-020 | 10 µg/ml | Live imaging |
|  | FITC anti-mouse IgM | JIR | 715-095-140 | 5 µg/ml | pH probe (FACS) |
|  | Donkey anti-mouse IgM AlexaFluor647-biotin | In-house | 715-605-140 + Thermo 21343 | 10 µg/ml | IF |
|  | Hen Egg Lysozyme (HEL) | Sigma-Aldrich | #L6876 | 10 µg/ml | ELISA |
|  | HEL-biotin | In-house | Sigma-Aldrich # L6876 + Thermo 21338 | 1-10 µg/ml | For DQ-Ova probe/FACS |
| Antibodies (IF) | Anti-Rab5 | CST | 3547 | 1:150 | IF |
|  | Anti-Rab6 | CST | 9625S | 1:200 | IF |
|  | Anti-Rab7 | Santa Cruz | Sc-376362 | 1:100 | IF |
|  | Anti-Rab9 | CST | 5118 | 1:150 | IF |
|  | Anti-Rab11 | CST | 5589 | 1:200 | IF |
|  | Anti-EEA1 | Santa Cruz | Sc-6415 | 1:50 | IF |
|  | Anti-LAMP1 | DSHB | 1D4B | 1:75 | IF |
|  | Anti-CathepsinS | LSbio | B2550 | 1:50 | IF |
|  | Anti-CathepsinS | Santa Cruz | sc-271619 | 1:50 | IF |
|  | Anti-MHCII | Santa Cruz | Sc-59322 | 1:50 | IF |
|  | Anti-MHCII-AF488 | In-house | Sc-59322 + Thermo A20000 | 1:50 | IF/Live imaging |
|  | Anti-PCM1-AF647 | Santa Cruz | Sc-398365 AF647 | 1:200 | IF |
|  | Donkey-anti-rabbit IgG (H+L) AlexaFluor 488/555/647 | Thermo | A21206, A31572, A31573 | 1:500 | IF |
|  | Donkey anti-goat IgG (H+L) AlexaFluor 488/555 | Thermo | A11055, A21432 | 1:500 | IF |
|  | Mouse anti-rat IgG Fcy Fragment Specific AlexaFluor 488/RRx/647 | JIR | 212-545-104<br>212-295-104<br>212-605-104 | 1:500 | IF |
|  | Goat-anti-mouse IgG Fcy subclass 1 AlexaFluor 488/RRx/647 | JIR | 115-545-205<br>115-295-205<br>115-605-205 | 1:500 | IF |
| ELISA | Anti-mouse IL-2 | Nordic Biosite | 503804 | 4 µg/ml | ELISA |
|  | Anti-mouse IL-2, biotin | Nordic Biosite | 503702 | 1 µg/ml | ELISA |
|  | ExtraAvidin-AP (alkaline phosphatase) | Sigma-Aldrich | E2636 | 1:5000 | ELISA |
|  | FAST pNPP substrate tablet | Sigma-Aldrich | N2770-5SET | - | ELISA |
| Other | LysoTracker Deep Red | Thermo | L12492 | 125 nM | Live imaging |
|  | DQ-OVA-biotin | In-house | Thermo D12053<br>+EZ-Link<br>Maleimide-PEG2-biotin<br>(Thermo<br>21901BID) | 1:10 | FACS |
|  | Fibronectin | Sigma | F4759-2MG | 4 µg/ml | IF |

